# Nitric oxide regulates metabolism in murine stress erythroid progenitors to promote recovery during inflammatory anemia

**DOI:** 10.1101/2023.03.11.532207

**Authors:** Baiye Ruan, Yuanting Chen, Sara Trimidal, Imhoi Koo, Fenghua Qian, Jingwei Cai, John Mcguigan, Molly A. Hall, Andrew D. Patterson, K. Sandeep Prabhu, Robert F. Paulson

## Abstract

Inflammation skews bone marrow hematopoiesis increasing the production of myeloid effector cells at the expense of steady-state erythropoiesis. A compensatory stress erythropoiesis response is induced to maintain homeostasis until inflammation is resolved. In contrast to steady-state erythroid progenitors, stress erythroid progenitors (SEPs) utilize signals induced by inflammatory stimuli. However, the mechanistic basis for this is not clear. Here we reveal a nitric oxide (NO)-dependent regulatory network underlying two stages of stress erythropoiesis, namely proliferation, and the transition to differentiation. In the proliferative stage, immature SEPs and cells in the niche increased expression of inducible nitric oxide synthase (*Nos2* or *iNOS*) to generate NO. Increased NO rewires SEP metabolism to increase anabolic pathways, which drive the biosynthesis of nucleotides, amino acids and other intermediates needed for cell division. This NO-dependent metabolism promotes cell proliferation while also inhibiting erythroid differentiation leading to the amplification of a large population of non-committed progenitors. The transition of these progenitors to differentiation is mediated by the activation of nuclear factor erythroid 2-related factor 2 (Nfe2l2 or Nrf2). Nrf2 acts as an anti-inflammatory regulator that decreases NO production, which removes the NO-dependent erythroid inhibition and allows for differentiation. These data provide a paradigm for how alterations in metabolism allow inflammatory signals to amplify immature progenitors prior to differentiation.

**Key points:**

1. Nitric-oxide (NO) dependent signaling favors an anabolic metabolism that promotes proliferation and inhibits differentiation.
2. Activation of Nfe2l2 (Nrf2) decreases NO production allowing erythroid differentiation.

## Introduction

Bone marrow erythropoiesis produces erythrocytes at a constant rate to maintain erythroid homeostasis at steady state. In contrast, infection and tissue damage induce inflammation, which increases myelopoiesis at the expense of steady-state erythropoiesis(^1–3^). To compensate for this loss of erythroid production, an alternative response pathway, stress erythropoiesis, is induced to produce erythrocytes until the inflammation is resolved and bone marrow erythropoiesis can resume(^4^). Stress erythropoiesis utilizes progenitor cells that are distinct from steady-state erythropoiesis(^5–7^). They are derived from short-term reconstituting hematopoietic stem cells (CD34^+^Kit^+^Sca1^+^Lin^−^) that migrate from murine bone marrow to spleen and liver, where these cells are specified as stress erythroid progenitors (SEPs) without proceeding through the CMP and MEP intermediates associated with steady state erythropoiesis(^5, 8^). Stress erythropoiesis uses a different strategy (^5, 9^), where immature SEPs rapidly proliferate but do not differentiate, generating a transient amplifying population of SEPs. This population consists of three sub-populations: the most immature Kit^+^Sca1^+^CD34^+^CD133^+^ cells, which give rise to Kit^+^Sca1^+^CD34^-^CD133^+^ cells, and the more mature Kit^+^Sca1^+^CD34^-^CD133^-^ cells(^7^). The expansion of the transient amplifying population ensures sufficient progenitors, so that once serum erythropoietin (Epo) levels increase, the synchronous differentiation of progenitors will generate enough erythrocytes to alleviate the anemia(^5, 7, 10^).

Rapidly proliferating cells like cancer cells are supported by profound metabolic changes that direct the utilization of glucose and glutamine to drive the biosynthesis of macromolecules necessary for cell growth and division(^11–16^). These changes include a preference for aerobic glycolysis rather than mitochondrial oxidative phosphorylation (OXPHOS). Glycolytic metabolites support cell proliferation by entering into the branched pathways for anabolism, e.g. pentose phosphate pathway (PPP) generates ribose-5 phosphate for nucleotide biosynthesis(^17, 18^), and serine/glycine biosynthetic pathway generates amino acids, lipids and metabolites for one-carbon metabolism(^19^). In addition, glutamine metabolism drives the production of cellular building blocks like nucleotides(^14, 15^). These same metabolic adaptations are also utilized to regulate cell function and lineage-specific differentiation. Glycolysis and glutaminolysis act in concert to increase the production of nucleotides to drive steady-state erythroid commitment(^20^). Lipopolysaccharide (LPS) activation of Toll-like receptor 4 (TLR4) shifts macrophage metabolism towards aerobic glycolysis, which promotes the production of pro-inflammatory cytokines and Nitric oxide (NO), which consequently drives the differentiation of M1 macrophages and maintains their metabolism(^21–27^). Previously we showed that pro-inflammatory signals in the niche increase growth-differentiation factor 15 (GDF15) expression(^9^). GDF15 regulates several key enzymes involved in glutamine and glucose metabolism which are required for SEP proliferation(^28^). These data suggest that the proliferation of SEPs is regulated by inflammation-induced metabolism. However, it remains unclear how inflammatory signaling alters metabolism, and how metabolic changes modulate proliferation and differentiation of SEPs.

Here, we show that immature SEPs and cells in the niche increase in NO production, which establishes a proliferative metabolism dominated by anabolic pathways that support cell proliferation. This NO-dependent metabolism also inhibits differentiation. The transition to differentiation leads to an activation of nuclear factor erythroid 2-related factor 2 (Nfe2l2 or Nrf2), which decreases NO production and removes erythroid inhibition. These data support a model where the same inflammatory signals that inhibit steady-state erythropoiesis, drive a compensatory stress erythropoiesis response that relies on an inflammatory metabolic profile to promote the expansion of early SEP populations needed to ensure efficient production of erythrocytes to maintain homeostasis.

## Material and methods

### Mice and cell culture

Wild-type C57BL/6J, B6.129P2-Nos2^tm1Lau^/J (Nos2-/-) Jax stock #002609 (^29^) and B6.129X1-Nfe2l2^tm1Ywk^/J(^30^) (Nrf2-/-) JAX stock #017009 mice were obtained from Jackson Laboratories. Age- and sex-matched mice were randomly assigned to different experimental groups. All experiments were performed according to protocols authorized by the Institutional Animal Care and Use Committee (IACUC) at the Pennsylvania State University. Murine and human stress erythropoiesis cultures were performed following a previous method(^7^). Detailed *in vitro* and *in vivo* experimental procedures are described in the supplemental methods.

### Statistical analysis

GraphPad Prism was used for statistical analysis. Data are presented as mean ± SD with individual data points displayed. The number of biological replicates and the statistical tests used for each experiment are detailed in the figure legends. Less than 0.05 of p value is considered as significant difference. n.s., p > 0.05; *, p < 0.05; **, p < 0.01; ***, p < 0.001.

### Data sharing statement

RNA-sequencing data have been deposited at NCBI GEO with accession number GSE190030. Metabolomics data have been deposited at NMDR (DOI: http://dx.doi.org/10.21228/M89402).

## Results

### Metabolism is rewired to increase anabolic biosynthesis for SEP proliferation

To analyze metabolism in proliferating SEPs, we used *in vitro* stress erythropoiesis cultures. Growing bone marrow cells in stress erythropoiesis expansion media (SEEM) recapitulates the expansion of SEPs observed *in vivo*(^7, 9, 10, 28, 31, 32^). SEPs rapidly expanded in SEEM showing an accumulation of Kit^+^Sca1^+^ cells over the 5-day culture (Figure 1A-B, S1A). In addition, the predominant progenitor population switched from Kit^+^Sca1^+^CD34^+^CD133^+^ (day 1 and 3) to Kit^+^Sca1^+^CD34^-^CD133^-^ cells at day 5. The latter population is more mature and highly proliferative(^28^).

**Figure 1.**
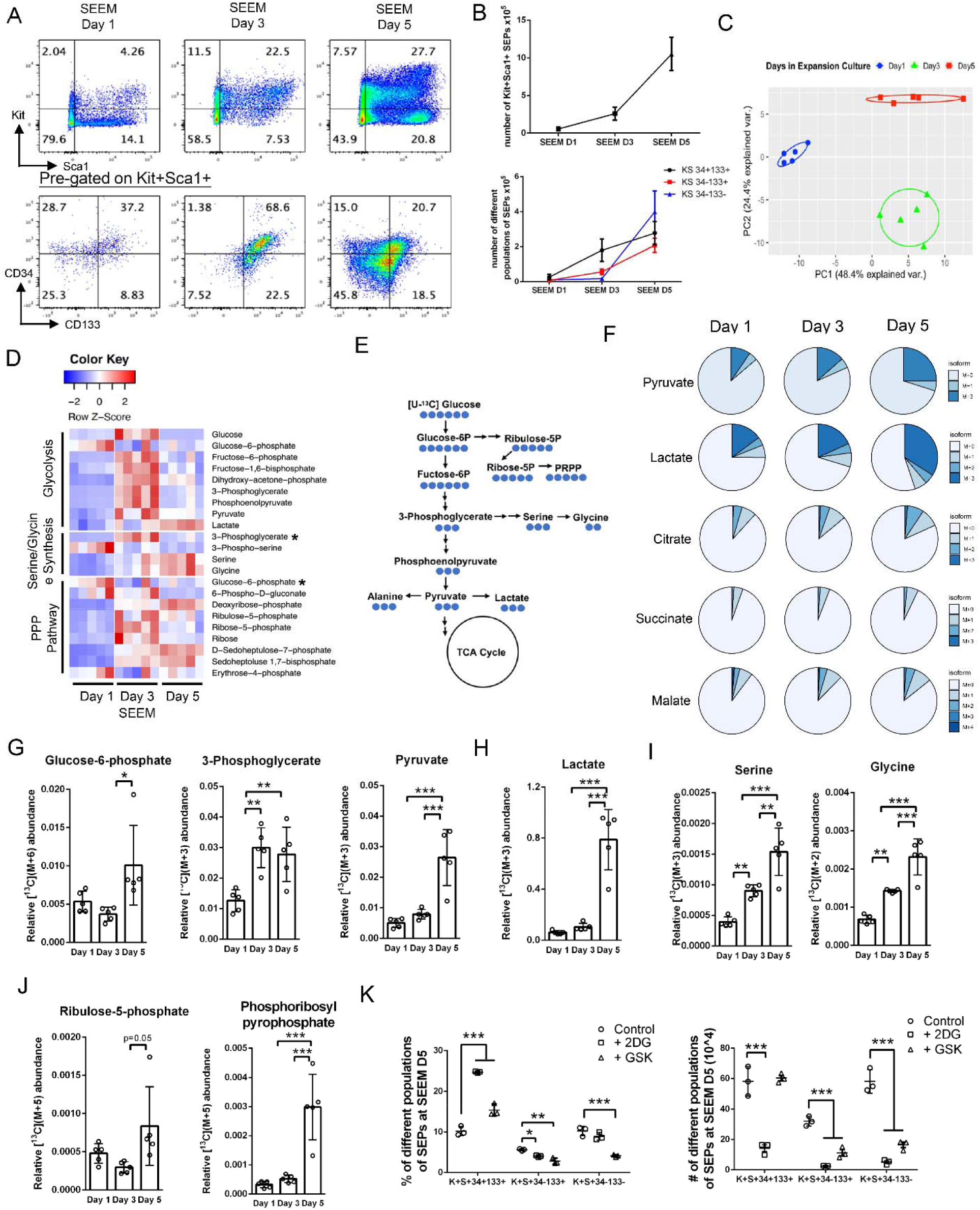
Increased glycolysis and diverging anabolic pathways in immature SEPs drive the biosynthesis of nucleotides and amino acids for rapid cell proliferation. (A-B) SEPs were isolated from SEEM cultures at indicated days for flow cytometry analysis. Cells were stained for viability followed by gating on Kit and Sca1. Pre-gated Kit+Sca1+ cells were then gated on CD34 and CD133. Representative flow cytometry plot (A) and quantification of absolute numbers of Kit^+^Sca1^+^ SEPs and indicated SEP populations (B) (n=5 per time point). (C-D) SEPs were isolated from SEEM cultures at indicated days for metabolomics analysis. PCA analysis of metabolomics profiling data (C). A heatmap depicting the abundance of metabolites in glycolysis, serine/glycine synthesis and PPP pathway extracted from SEPs from SEEM cultures over time. Metabolites involved in more than one pathway are marked by *. Color represents row-wise scaled z-score of metabolite abundance (D) (n=5 per time points). (E-J) SEPs isolated from regular SEEM on day 1, 3 and 5 were re-cultured in SEEM supplemented with 25 mM U-[^13^C]-Glucose. After another 24 hrs, SEPs were collected for metabolic profiling. Schematic for metabolism of [U-^13^C] glucose (E). ^13^C-isotope labeling distribution pattern of pyruvate, lactate and TCA cycle intermediates (F). Relative abundance of ^13^C-isotope labeling of glycolytic intermediates (G), lactate (H), serine and glycine (I), and PPP metabolites (J) (n=5 per time point, one-way ANOVA/Holm-Sidak). (K) SEEM cultures were treated with vehicle, 1 mM 2DG or 13 nM GSK at day 3 for 48 hrs. Flow cytometry analysis of percentages (left) and absolute numbers (right) of indicated SEP populations (n=3 per group, one-way ANOVA/Dunnett’s). Data represent mean ± SD. * p < 0.05, ** p < 0.01, *** p < 0.001.

Metabolites were extracted from bulk SEPs on days 1, 3 and 5 of cultures and analyzed by LC-MS analysis. Principle component analysis showed that intracellular metabolic pattern changed over culture time and were distinct as SEPs matured and proliferated (Figure 1C). Day-3 SEPs, which were predominantly Kit^+^Sca1^+^ CD34^+^CD133^+^ cells, showed higher levels of metabolites in glycolysis, PPP and serine/glycine biosynthetic pathway (Figure 1D), suggesting that the proliferating progenitors actively engage glucose metabolism to increase glycolytic and anabolic activities to support cell division. Consistently, we found a pronounced increase of metabolites in amino acid and nucleotide metabolism (Figure S1B). Day-5 SEPs maintained high accumulation of lactate, but the increase in other glycolytic metabolites was less, which suggested that by day 5 nutrients might be limiting in our cultures (Figure 1D). Despite the decrease in glycolysis, we observed a notable decrease in 3-phosphoglycerate (3PG) and an increase in serine and glycine (Figure 1D), which suggested that the glycolytic intermediate 3PG was utilized for serine/glycine biosynthesis. To further examine metabolism in SEPs, we performed metabolomic analysis on SEPs isolated from the spleen on days 0, 1 and 3 after phenylhydrazine (PHZ) treatment. Kit+ SEPs were isolated from spleen by magnetic bead sorting and sorted cells showed increased enrichment of SEP markers (Figure S1C). Kit+ SEPs were processed as described above for in vitro cultures. This analysis shows that glycolysis increases over the 3 days of recovery. The increase in glycolytic metabolites was not uniform. On day 1 after PHZ, early metabolites (glucose 6-phosphate, Fructose 6-phosphate and glyceraldehyde 3-phosphate) in glycolysis were increased, which corresponded to increased PPP metabolites, while on day 3 later glycolytic metabolites are increased which paralleled the increase in nucleotides and amino acid production (Figure S1D). Overall, the metabolomic data from in vivo SEPs was similar to that observed in vitro.

To examine glucose utilization, we measured the intracellular accumulation of ^13^C-labeled metabolites after supplementing day 1, 3 and 5 SEP cultures with uniformly labeled U-[^13^C]-glucose (Figure 1E). Proliferating SEPs actively use glucose for glycolysis, as indicated by the increased isotopic enrichment of ^13^C-glucose-derived glycolytic intermediates at later time points when compared to day 1 progenitors (Figure 1F-H, S2A). Labeling distribution pattern indicated that pyruvate transfers carbon to lactate, whereas it has limited entry into TCA cycle (Figure 1F). Consistently, the increase of labeling of TCA cycle intermediates was not as great as the increase in lactate labeling (Figure S2B), suggesting that proliferating SEPs preferentially use glucose-derived pyruvate for glycolysis over OXPHOS. Glucose-6-phosphate (G6P) can be shunted from glycolysis into PPP by generating ribulose-5-phosphate (Ru5P). Day-5 SEPs showed elevated enrichment of M+5 isotopologues of Ru5P, and the pattern of which was coupled with M+6 G6P. Additionally, we observed increased labeling of phosphoribosyl pyrophosphate (PRPP), a metabolite derived from Ru5P that acts as a precursor for increased purine and pyrimidine biosynthesis (Figure 1J, S2C). Day-3 SEPs showed greater 3PG, serine and glycine labeling than day 1, whereas little to no difference was detected in M+3 pyruvate and lactate, indicating that 3PG is diverted from glycolysis to the serine/glycine biosynthesis pathway (Figure 1G-I) and pyruvate is diverted to generate alanine (figure S2D). Blocking glycolysis using 2-Deoxy-D-glucose (2-DG) significantly impaired the proliferation of Kit+Sca1+ SEPs (Figure S2E). All three populations were affected by 2-DG. In contrast, the lactate dehydrogenase (Ldh) inhibitor GSK 2837808A(^33^) (GSK) also impaired SEP proliferation, but to a lesser extent (Figure S2E). However, GSK did not affect the number of immature Kit^+^Sca1^+^CD34^+^CD133^+^ progenitors, while decreasing the number of Kit^+^Sca1^+^CD34^-^CD133^-^ cells (Figure 1K), the latter of which proliferate faster and have increased expression of genes involved in cell cycle and anabolic biosynthesis(^28^). This difference in response most likely reflects the fact that 2DG affects the entire glycolytic pathway as well as the anabolic pathways that utilize glycolytic metabolites.

These data demonstrate that the early proliferating SEPs actively utilize glycolysis and shunt glycolytic intermediates into its branched pathways generating anabolic intermediates to promote cell proliferation.

### Glutaminolysis increases AASS flux to fuel iNOS-dependent NO production in proliferating SEPs

The metabolic profiles we observed in early SEPs are like other highly proliferative cells such as tumor and activated immune cells, where iNOS signaling acts as a key metabolic regulator(^21, 24, 34^). Stress erythropoiesis is promoted by inflammatory stimuli, zymosan and LPS, and pro-inflammatory cytokines, TNF-α and IL-1β . Immature SEPs can directly respond to these inflammatory signals, as indicated by a profound expression of their receptors (Figure S3A).

Expression of *Nos2* (iNOS) elevated in SEEM cultures over time, whereas the other two NO synthases, *Nos1* and *Nos3*, were expressed at barely detectable levels (Figure 2A). We observed increased NO levels in proliferating SEPs, however, the increase was disrupted when SEEM cultures were treated with 1400w, a selective inhibitor for iNOS (Figure 2B, S3B-C). These results reveal that proliferating SEPs induce iNOS and are one source of NO in the niche. To determine the nutrient sources for NO synthesis, SEEM cultures were treated with 2-DG to block glucose metabolism, or 6-diazo-5-oxo-L-norleucine (DON)(^35^) to block glutamine metabolism. Addition of 2-DG had no effect on NO MFI levels, the NO+ population in 2DG treated cultures was broader than control. In contrast, DON resulted in a sharp decrease, which was further decreased when DON and 2-DG were added together (Figure 2C). Furthermore, treatment of DON for as little as 3h was enough to attenuate NO levels in SEPs (Figure S3D). Glutamine is converted to glutamate by glutaminase (Gls), which enters AASS via the enzyme aspartate aminotransferase (Got1). The flux through AASS replenishes TCA cycle and at same time provides substrates for NO synthesis(^22^). The proliferating SEPs displayed increased levels of metabolites and mRNA expression of key enzymes associated with AASS (Figure 2D-E). This result is consistent with our previous data showing that *Gls* and *Got1* are more highly expressed in rapidly proliferating Kit^+^Sca1^+^CD34^-^CD133^-^ SEPs when compared to the immature Kit^+^Sca1^+^CD34^+^CD133^+^ cells^27^. The addition of Gls inhibitor, DON, or Got1 inhibitor, aminooxyacetic acid(^36^) (AOA), diminished all identified SEP populations in the expansion cultures (Figure S3E-H). These data support the use of glutamine metabolism to increase the flux through AASS to support NO synthesis.

**Figure 2.**
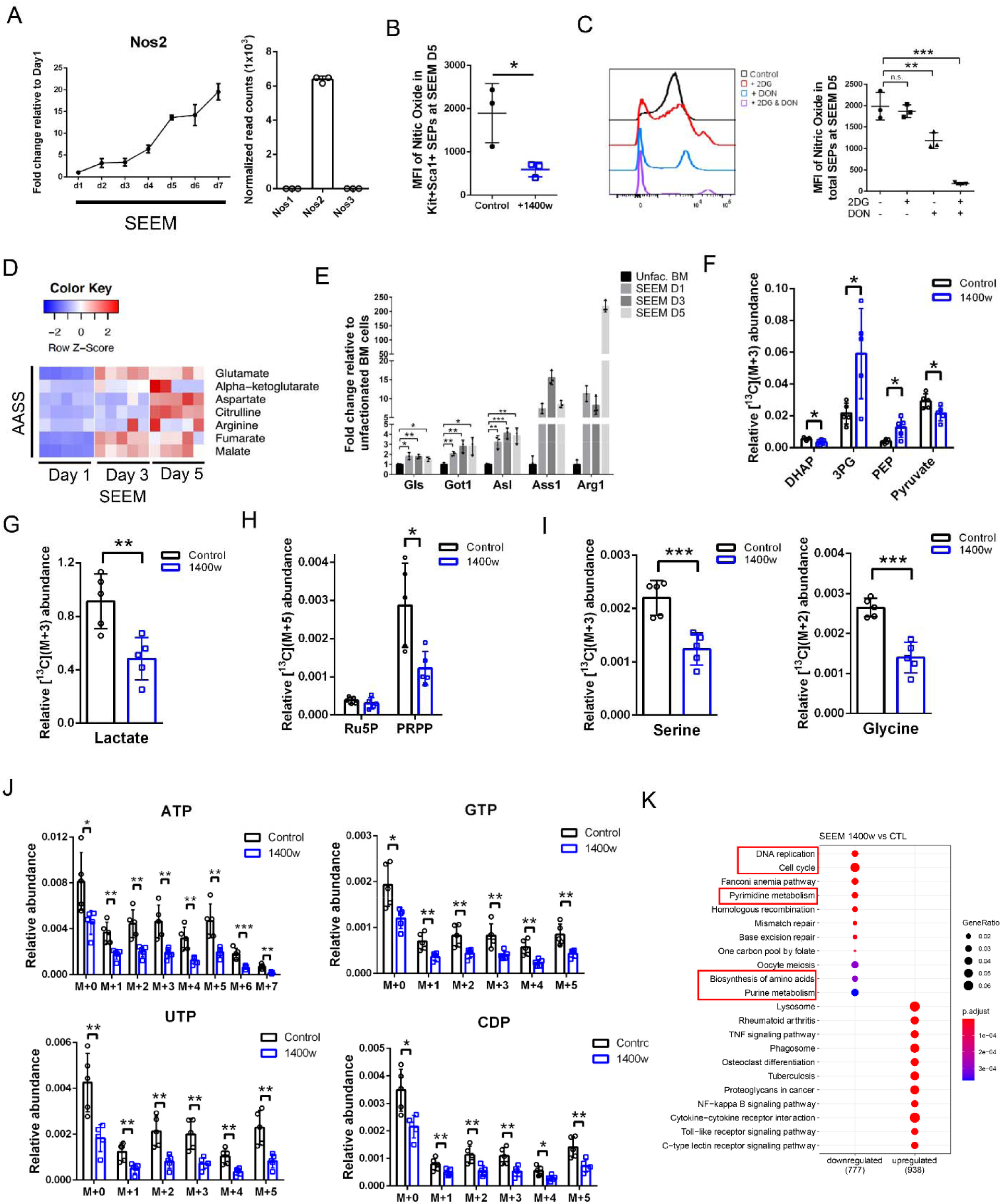
Glutaminolysis fuels iNOS-dependent NO production to maintain a proliferative metabolic profile in immature SEPs. (A) (left) qRT-PCR analysis of *Nos2* expression in SEPs grown in SEEM over time (n=3 per time point). (right) RNA-seq data showing *Nos1, Nos2 and Nos3* expression in SEPs at SEEM day 5 (n=3). (B) The intracellular levels of NO in Kit^+^Sca1^+^ SEPs treated ± 10 M μ1400w quantified by mean fluorescence intensity (MFI) of DAF-FM DA staining (n=3 per group, unpaired t test). (C) Quantification of NO levels in SEPs from SEEM cultures treated with vehicle, 1 mM 2DG or μM DON alone, or in combination at day 3 for 48 hrs. Representative flow cytometric plot (left) and quantification of MFI (right) of NO staining (n=3 per group, one-way ANOVA/Dunnett’s). (D) A heatmap depicting the abundance of AASS metabolites in SEPs isolated from SEEM over time (n=5 per time point). (E) qRT-PCR analysis of expression of genes related to AASS from unfractionated BM cells and SEPs on indicated days of SEEM cultures (n=3 per group, multiple t test/Holm-Sidak). (F-J) SEEM cultures were treated with 10 μM 1400w or vehicle at day 3 for 48 hrs, and then cells were re-cultured in SEEM with U-[13C]-Glucose for 24 hrs. Relative abundance of 13C711 isotope labeling of selected glycolytic metabolites (F), lactate (G), PPP metabolites (H), serine and glycine (I), and metabolites in nucleotide synthesis (J) (n=5 per group, unpaired t test). (K) RNA-seq analysis comparing SEPs treated ± 10 μM 1400w at SEEM day 3 for 48 hrs. Top enriched KEGG pathways identified via overrepresentation analysis of downregulated DEGs (FDR < 0.05, fold change (FC) of 1400w/CTL < -1.5) and upregulated DEGs (FDR<0.05, FC > 1.5). Circle size represents gene ratio and color represents BH-adjusted p value. Red box highlights pathways related to cell proliferation and anabolic pathways (n=3 per group). Data represent mean ± SD. n.s. p > 0.05, * p < 0.05, ** p < 0.01, *** p < 0.001.

### iNOS-derived NO is required to maintain the proliferating SEP metabolism

To examine the role of iNOS in SEP metabolism, we studied incorporation of U-[^13^C]-glucose into metabolites extracted from SEPs after treating cultures with or without the Nos2 specific inhibitor, 1400w(^37^). The inhibition of iNOS led to a glycolytic break, where we observed reduced ^13^C-glucose-labeled pyruvate and an accumulation of upstream metabolites PEP and 3PG (Figure 2F), indicating an inhibition of pyruvate kinase-mediated conversion of PEP to pyruvate. However, RNA-seq data showed that the suppression of pyruvate kinase activity did not occur at transcriptional level (Figure S4A). The major form of PK expressed in SEPs is the PKM2 isoform of PKM, which is known to be regulated by nitrosylation(^38–40^) (Figure S4B). The decrease in labeled pyruvate was mirrored by decreased levels of labeling in downstream metabolites lactate (Figure 2G) and alanine (Figure S4C). In contrast, TCA cycle intermediates (except for malate) were only slightly affected (Figure S4D). Despite increased levels of 3PG and G6P, the levels of labeled serine and glycine were decreased as were the levels of Ru5P and PRPP as well as downstream nucleotides, which showed that shunting of metabolites to anabolic pathways was impaired (Figure 2F, H-J, S4E). Consistent with the results from isotope tracing experiment, RNA-seq analysis showed that iNOS inhibition suppressed the expression of genes involved in anabolic pathways, including biosynthesis of amino acids, pyrimidine and purine metabolism (Figure 2K, S4F). In particular, the expression of Pfkfbp3 was increased in cells treated with 1400W, while untreated control cells expressed higher levels of Pfkfbp4.

Pfkfbp3 exhibits a higher kinase activity compared to its phosphatase activity and drives metabolites through the glycolytic pathway. In contrast, Pfkfbp4 has a higher phosphatase activity, which drives metabolites back to glucose 6-phosphate so it can enter the PPP and drive nucleotide metabolism(^41^) (Figure S4G). Similarly, in vivo metabolomic analysis Nos2-/-SEPs isolated on day 3 after PHZ treatment showed defects in glycolysis, PPP, amino acid and nucleotide biosynthesis (Figure S4H). Overall, these observations indicate that stress erythropoiesis induces iNOS-dependent synthesis of NO to maintain the active glycolysis and anabolic biosynthesis in early proliferating progenitors.

### iNOS-dependent synthesis of NO promotes SEP proliferation

RNA-seq data showed that inhibiting iNOS with 1400w resulted in the suppression of a spectrum of biological processes associated with cell cycle, DNA replication and mitosis (Figure 2K, 3A, S5A-B). Consistent with these changes in gene expression, we observed fewer overall SEPs when cultures were treated with 1400w (Figure 3B). However, this decrease was due to fewer more mature, rapidly proliferating Kit^+^Sca1^+^CD34^-^CD133^+^ and Kit^+^Sca1^+^CD34^-^CD133^-^ progenitor populations (Figure 3C). The block of SEP proliferation was reproduced using another iNOS inhibitor L-NIL(^42^), or iNOS knockouts in which additional treatment of 1400w or a non-specific NOS inhibitor, LNNA(^43^), did not exacerbate the decrease of SEPs, which demonstrates that Nos2 is the primary source of NO in the in vitro cultures (Figure 3D, S5C-D). However, when Nos2-/-cells were co-cultured with control cells, control cells were able to rescue the defect in proliferation, which shows that SEPs and cells in the niche make NO (Figure S5E). We further show that in vivo metabolomic analysis of Nos2-/-SEPs isolated on day 3 after PHZ treatment showed defects in glycolysis, PPP, amino acid and nucleotide biosynthesis (Figure S4H).

**Figure 3.**
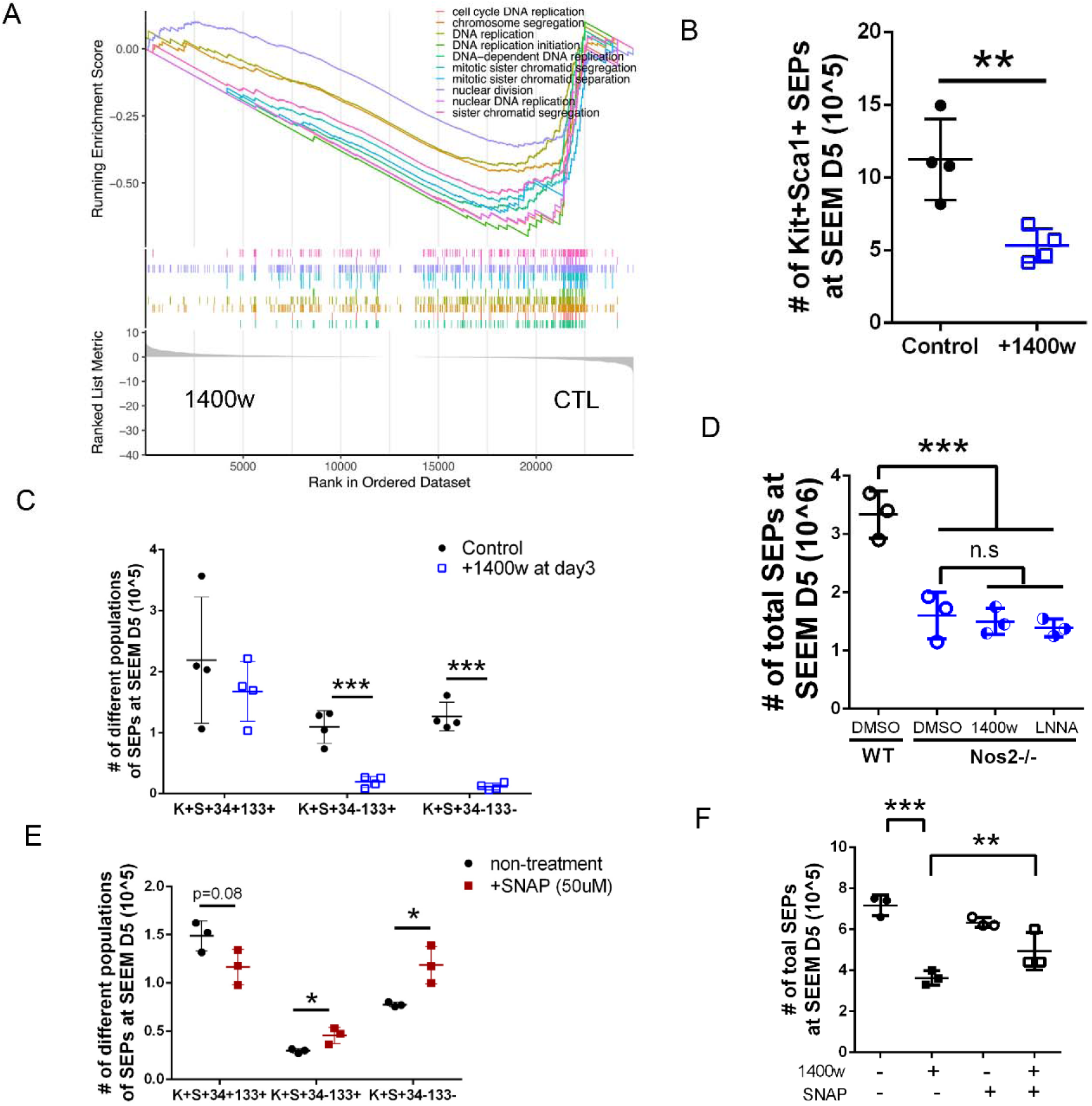
iNOS-derived NO is essential for *in vitro* expansion of immature SEPs. (A) GSEA analysis of cell cycle-related GO terms between 1400w-vs CTL-treated SEPs as described in Fig 2K (n=3 per group). (B-C) Kit+Sca1+ SEP cell counts (B), and flow cytometry analysis of the absolute numbers of indicated SEP populations (C) in cultures treated ± 10 μ test). (D) SEP cell counts in WT, or Nos2-/-SEEM cultures treated with vehicle, 10 μM LNNA (n=3 per group, one-way ANOVA/Dunnett’s). (E) The numbers of indicated populations of SEPs in SEEM cultures treated ± 50 μM SNAP (n=3 per group, unpaired t test). (F) SEP cell counts in SEEM cultures treated with 10 μM 1400w or 10 μM SNAP alone, or in combination for 24 hrs (n=3 per group, one-way ANOVA/Tukey’s). Data represent mean ± SD. n.s. p > 0.05, * p < 0.05, ** p < 0.01, *** p < 0.001.

In contrast to the effects of iNOS inhibition, increasing NO levels by treating cultures with the NO donor SNAP(^44^) accelerated the transition of SEPs from early Kit^+^Sca1^+^CD34^+^CD133^+^ to faster proliferating Kit^+^Sca1^+^CD34^-^CD133^-^ cells (Figure 3E). However, higher concentrations of SNAP, >50μ

M SNAP, decreased proliferation (Figure S5F). In addition, SNAP ameliorated the decrease of SEP numbers caused by iNOS inhibition (Figure 3F, S5G), which demonstrated that SEP proliferation relies on iNOS-dependent production of NO. To examine the role of iNOS signaling in SEP expansion *in vivo*, we induced acute hemolytic anemia in C57BL/6 wild-type (WT) and Nos2-/-mice by PHZ injection. Untreated Nos2-/-mice have similar numbers of spleen cells and SEPs in their spleens when compared to controls (Figure S5H). Compared to WT controls, Nos2-/-mice exhibited a defect in the expansion of cell populations in the spleen as demonstrated by decreased spleen weight and spleen cellularity (Figure 4A-C, S5I-J). We also observed a delayed and defective expansion of the faster proliferating Kit^+^Sca1^+^CD34^-^ CD133^+^ and Kit^+^Sca1^+^CD34^-^CD133^-^ SEP populations (Figure 4C, S5K). Although Nos2-/-mice exhibited a more severe anemia with decreased hematocrit, RBC numbers and hemoglobin values, they recovered, which may be in part due to the increased expression of *Nos1* in the spleens of Nos2-/-mice, which could compensate for the loss of Nos2-dependent NO production (Figure 4D-F, S5I). In addition, Nos2-/-spleens had increased infiltrating monocytes which could produce other inflammatory signals to compensate for SEP expansion in vivo (Figure S5L). We observed similar defects in recovery of Nos2-/-mice using a model of inflammatory anemia induced by heat-killed *Brucella abortus* (HKBA)(^45^). Nos2-/-mice were slower to recover from HKBA treatment and generated fewer stress BFU-E at days 8, 12 and 16 and HKBA treatment (Figure 4G-H). Analysis of normal erythropoiesis indicated that iNOS is not required for bone marrow erythropoiesis (Figure S5M-O, 5C). Collectively, these data demonstrate that iNOS-derived NO promotes the proliferation of SEPs *in vitro* and *in vivo* during the recovery from anemia.

**Figure 4.**
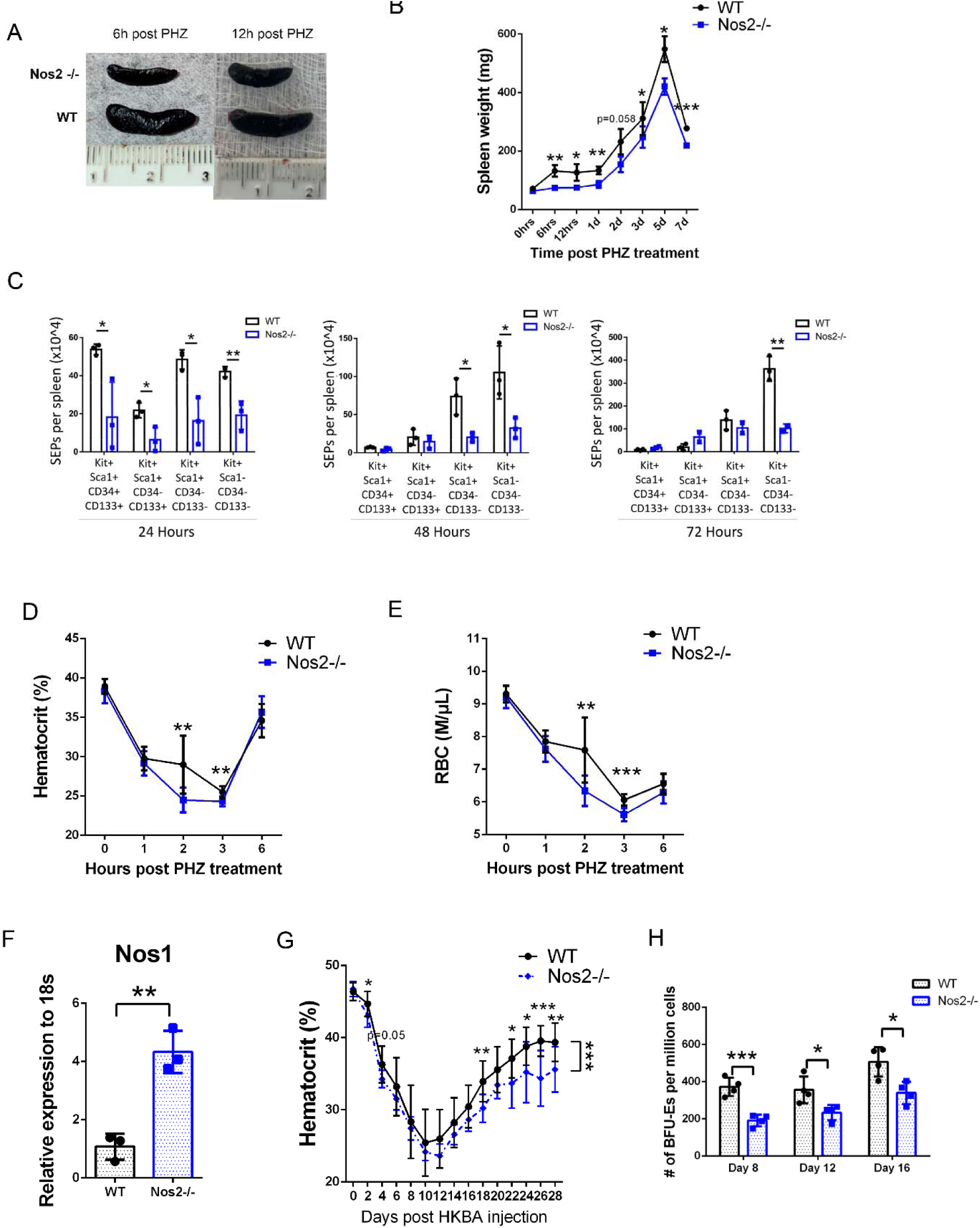
Stress erythropoiesis utilizes NO to expand progenitors for *in vivo* erythroid regeneration. (A-E) Age- and sex-matched WT and Nos2-/-mice were injected intraperitoneally with 100 mg/kg body weight of PHZ. Time course analysis of recovery from PHZ-induced hemolytic anemia. Spleen size (A), spleen weight (B), flow cytometry analysis of the absolute numbers of indicated SEP populations at 24 (left) , 48 (center) and 72 (right) hours after PHZ treatment (C), hematocrit (D), RBC counts (E) and qRT-PCR detection of *Nos1* mRNA abundance in splenocytes (F) of WT and Nos2-/-mice at indicated time points post PHZ injection (n=3 (B-C, F); n=7-8 (D-E), unpaired t test). (G-H) Age- and sex-matched mice were administered intraperitoneally with HKBA at 5 x 10^8^ particles/mouse. Time course analysis of recovery from HKBA-induced inflammatory anemia. Hematocrit (G) and stress BFU-E quantification (H) in WT and Nos2-/-mice on indicated days after injection with HKBA (n=12 in WT, n=8 in Nos2-/-, repeated measures two-way ANOVA followed by unpaired t test (G); n=3, unpaired t test (H)). Data represent mean ± SD. * p < 0.05, ** p < 0.01, *** p < 0.001.

### Inhibition of NO production enables activation of the erythroid gene expression program during differentiation

Epo signaling drives the transition from proliferating SEP to committed erythroid progenitor(^5, 7, 10^). Switching *in vitro* stress erythropoiesis cultures to differentiation media (SEDM) resulted in an increase in erythroid gene expression (Figure S6A) and a decrease in *Nos2* expression (Figure 5A). We observed a similar response *in vivo*, where *Nos2* expression increased in the first 24 hours, but rapidly decreased back to base level by 48 hours after PHZ treatment (Figure 5B and S5O). The appearance of stress BFU-E in the spleen follows this decrease(^46^). Levels of NO in SEPs in the spleen also increase over the first 24 hours after which they rapidly decline as progenitors differentiate (Figure 5C).

**Figure 5.**
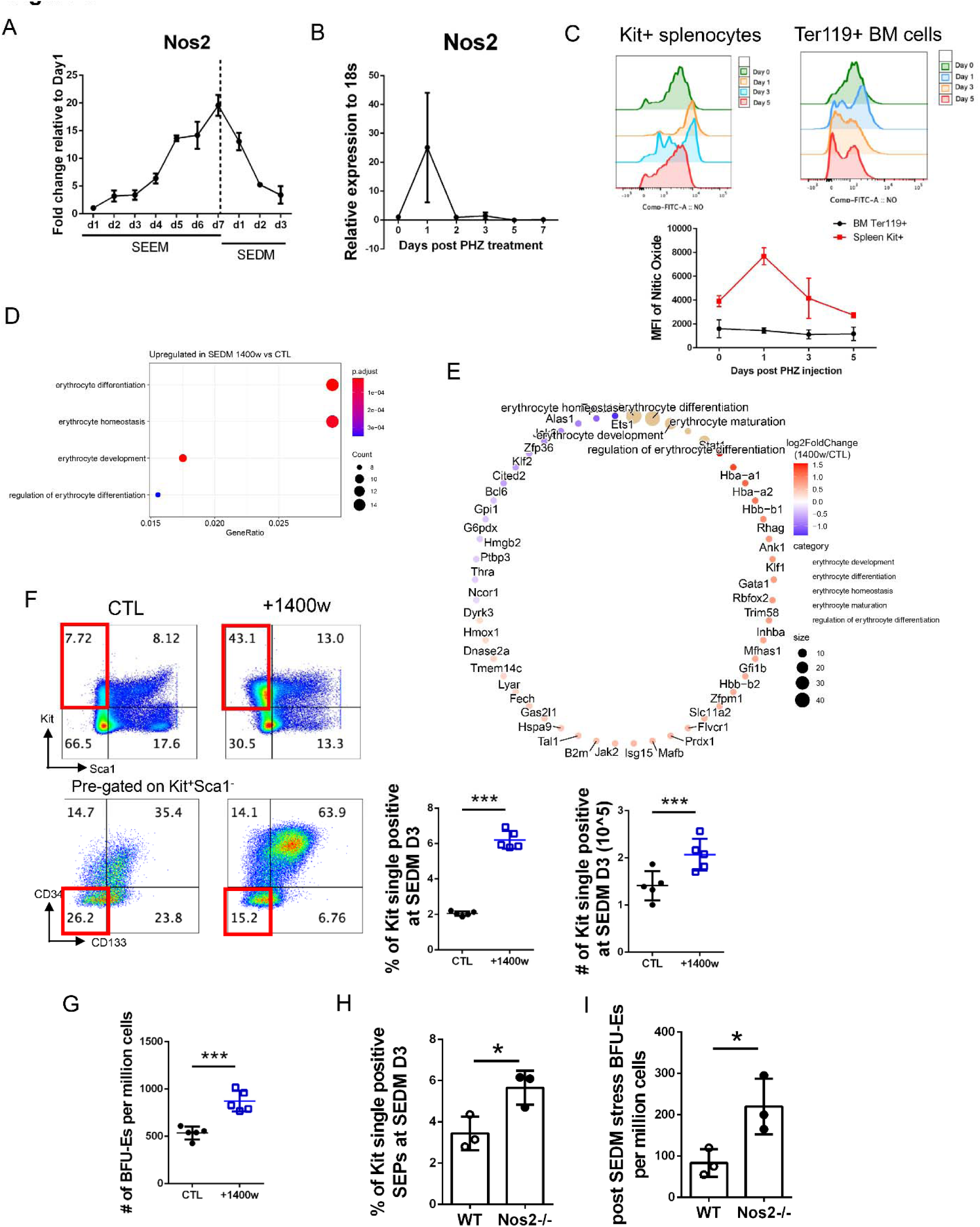
Inhibition of NO production enables the activation of the erythroid gene expression program during differentiation. (A) qRT-PCR analysis of *Nos2* expression at indicated days of SEEM and SEDM cultures (n=3 per time point, performed simultaneously with Fig 2A). (B) Splenocytes were harvested at indicated time points post PHZ treatment, followed by qRT-PCR analysis of *Nos2* mRNA abundance (n=3 per time point). (C) Quantification of NO levels in Kit+ splenocytes and Ter119+ BM cells (n=3 per time point). (D-E) After a 5-day culture in SEEM, SEPs were switched to SEDM and were treated with μM 1400w for 3 days. RNA-seq result assessed by overrepresentation test of upregulated DEGs by 1400w (FDR < 0.05, FC of SEDM 1400w/CTL > 1.5), depicting enrichment of GO terms related to erythropoiesis. Circle size represents the numbers of genes in each GO term and color represents BH-adjusted p value (D). Network analysis of DEGs (FDR < 0.05, |FC| > 1.2) in erythroid-associated pathways with color representing log2FC of SEDM 1400w/CTL (E) (n=3 per group). (F-G) SEDM cultures were treated with vehicle or 1400w as described in (D-E). Representative flow cytometry plot showing Kit+Sca1-CD34-CD133-cells (left), and quantification of percentages (middle) and absolute numbers (right) of Kit^+^Sca1^-^CD34^-^CD133^-^ differentiating SEPs (F). Quantification of stress BFU-Es per million cells by colony assay (G) (n=5 per group, unpaired t test). (H-I) SEPs in WT and Nos2-/-SEDM cultures were analyzed for the percentages of Kit^+^Sca1^-^ CD34^-^CD133^-^ SEPs (H) and stress BFU-E colonies per million cells (I) (n=3 per group, unpaired t test). Data represent mean ± SD. n.s. p > 0.05, * p < 0.05, ** p < 0.01, *** p < 0.001.

Inhibiting iNOS activity with 1400w in proliferating cells grown in SEEM resulted in increased expression of erythroid-specific genes (Figure S6B-C). While treatment of differentiation cultures (SEDM) with 1400w increased erythroid commitment over and above what was observed with Epo alone (Figure 5D-E, S6D). 1400w-treated cultures generated more mature Kit^+^Sca1^-^CD34^-^CD133^-^ SEPs and generated a significantly higher frequency of stress BFU-E (Figure 5F-G). The activation of SEP differentiation was reproduced by treating SEDM with L-NIL, or when Nos2-/-cultures were switched into SEDM media (Figure 5H-I, S6E-F).

### Nrf2 promotes differentiation by alleviating NO mediated suppression of erythroid differentiation

Inflammation drives NO production in phagocytes, which in turn induces a Nrf2-dependent anti-inflammatory feedback loop to suppress NO production(^47^). We reasoned that the decrease of NO production in differentiating progenitors is also mediated by Nrf2. Our *in vitro* analysis showed that the expression of the Nrf2 target gene, *Nqo1* increased when cells were switched into differentiation media, conversely, *Nos2* expression decreased (Figure 6A, 5A). During the recovery from PHZ-induced anemia *In vivo*, we observed that Nrf2 target genes *Nqo1* and *Gclm* increased after PHZ treatment, while *Nos2* decreased in the spleen (Figure 6B). RNA-seq analysis comparing gene expression of Nrf2-/- and control SEPs isolated from SEDM cultures showed that Nrf2-/-SEPs expressed higher levels of pro-inflammatory makers including *Il1b*, *Ptgs2* and *Nos2* (Figure 6C). In contrast, the genes downregulated by Nrf2 deficiency were involved in erythroid differentiation and hemoglobin biosynthesis (Figure 6C, S7A). Heatmap analysis using these genes as an erythroid program signature indicated that Nrf2 mutation prevents the increase in erythroid genes, while inhibition of iNOS with 1400W super induces the expression of these genes (Figure 6D). These data suggest that Nrf2 acts to counteract NO-dependent inhibition of the erythroid transcriptional program.

**Figure 6.**
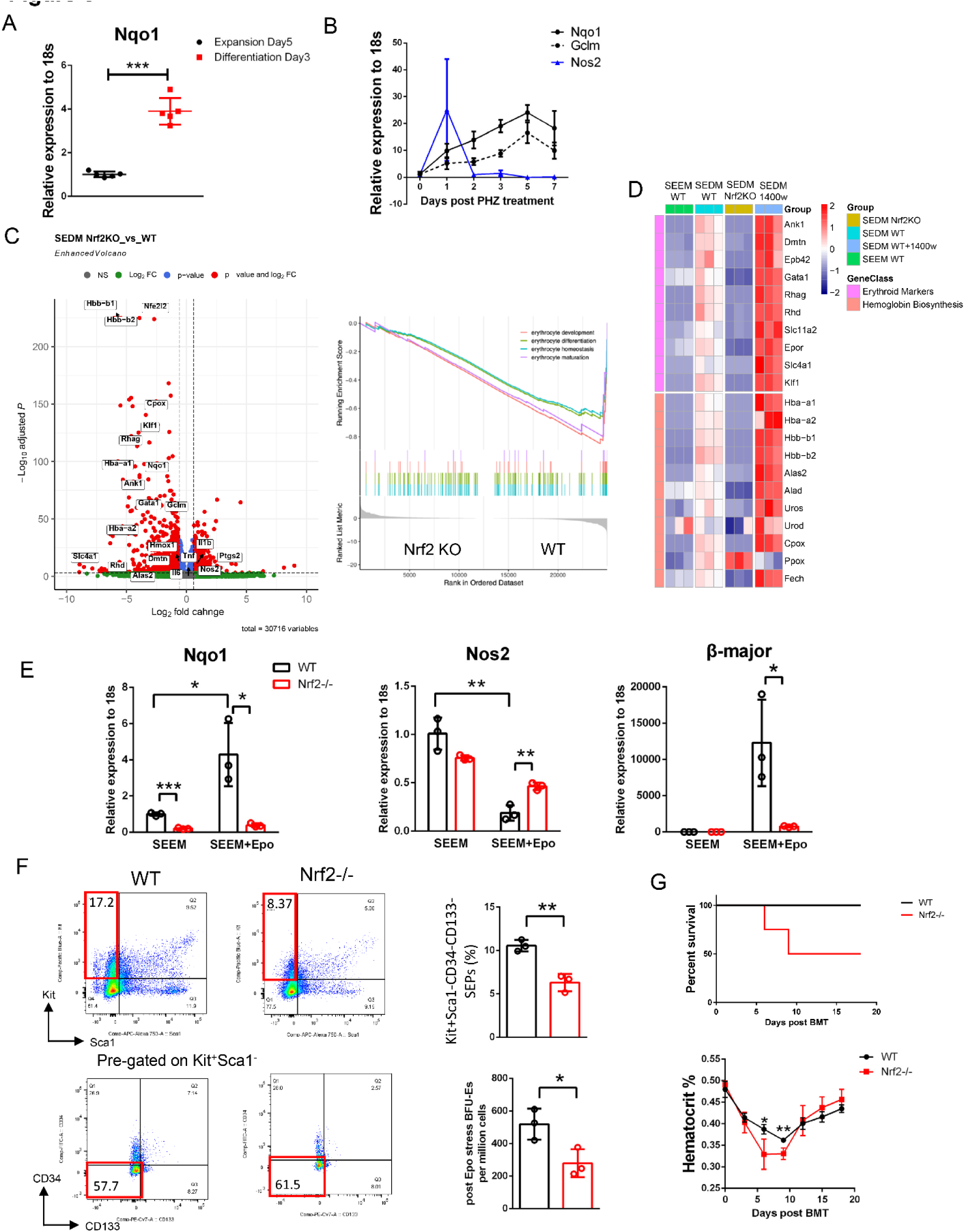
Activation of Nrf2 during differentiation counteracts NO-dependent erythroid inhibition. (A) qRT-PCR detection of *Nqo1* mRNA abundance between SEEM Day5 and SEDM Day3 SEPs (n=5 per group, unpaired t test). (B) qRT-PCR detection of *Nqo1*, *Gclm* and *Nos2* mRNA abundance from WT mice splenocytes at indicated time points after PHZ treatment (n=3 per condition). (C) Volcano plot of RNA-seq data comparing DEGs between WT and Nrf2-/-SEPs in SEDM cultures (left) and GSEA analysis of erythroid-associated GO terms (right) (n=3 per group). (D) Heatmap analysis of the progenitor RNA-seq profiles from the indicated conditions, depicting the expression levels of selected genes involved in erythroid differentiation and hemoglobin biosynthesis. Color bar represents row-wise scaled z-score (n=3 per group). (E-F) After a 5-day culture in SEEM, WT and Nrf2-/-SEPs were switched to Epo-containing SEEM for another 5-day culture in normoxia (20% O_2_). SEPs were harvested at indicated time points for qRT-PCR analysis of *Nqo1*, *Nos2* and β flow cytometry plot (left) and quantification of percentages of Kit^+^Sca1^-^CD34^-^CD133^-^ differentiating SEPs (top right), and quantification of frequency of stress BFU-Es by colony assay (bottom right) post Epo-containing SEEM culture (F) (n=3 per condition, unpaired t test). (G) Analysis of erythroid short-term radioprotection after BMT. WT or Nrf2-/-recipient mice were transplanted with 5 x 10^5^ donor BM cells derived from the same mouse strain (WT to WT; Nrf2-/-to Nrf2-/-). Analysis of survival (top) and hematocrit (bottom) during the recovery from BMT (n=4, unpaired t test). Data represent mean ± SD. * p < 0.05, ** p < 0.01, *** p < 0.001.

We verified the role of Nrf2 in differentiation using a Nrf2 activator, tert-butylhydroquinone(^48^) (tBHQ), which disrupts the interaction between Nrf2 and Keap1. tBHQ-treated differentiation cultures generated more mature Kit+Sca1-CD34-CD133-SEPs and stress BFU-Es, as well as expressed higher levels of erythroid-specific genes (Figure S7B-D). Differentiation cultures are incubated under hypoxia (2% O_2_) to potentiate Epo-induced transition to differentiation. To eliminate the impact of hypoxia, we performed WT and Nrf2-/-differentiation cultures in ambient air (20% O_2_). Epo was sufficient to activate Nrf2 without hypoxia, which in turn inhibited *Nos2* expression (Figure 6E). The defects of erythroid commitment in Nrf2 mutants were reproduced in cultures under normoxia (Figure 6F, S7E). We also observed that transplant of Nrf2-/-bone marrow into irradiated recipients resulted in slower erythroid recovery and decreased survival supporting a role for Nrf2-dependent regulation of stress erythropiesis *in vivo* (Figure 6G).

To determine whether Nrf2 is directly involved in increasing erythroid gene expression or it activates the erythroid program by blocking iNOS, we isolated SEPs from control and Nrf2-/-SEDM cultures treated with and without 1400w. We found higher levels of NO in Nrf2-/-SEPs, whereas NO levels returned to control levels in Nrf2-/-cultures treated with 1400w, which indicated that Nrf2 activation impairs iNOS-dependent NO synthesis (Figure 7A). 1400w treatment had no effect on the expression of Nrf2 target genes (Figure 7B). We next evaluated the erythroid differentiation of SEPs by analyzing the generation of Kit single positive SEPs, stress BFU-Es, and the expression of erythroid-associated genes. Although we observed severe defects in SEP differentiation resulting from Nrf2 mutation, they were completely rescued by the inhibition of iNOS by 1400w (Figure 7C-F). Our data demonstrate that Nrf2 promotes SEP differentiation by inhibiting the expression of *Nos2*, which relieves NO-mediated erythroid inhibition.

**Figure 7.**
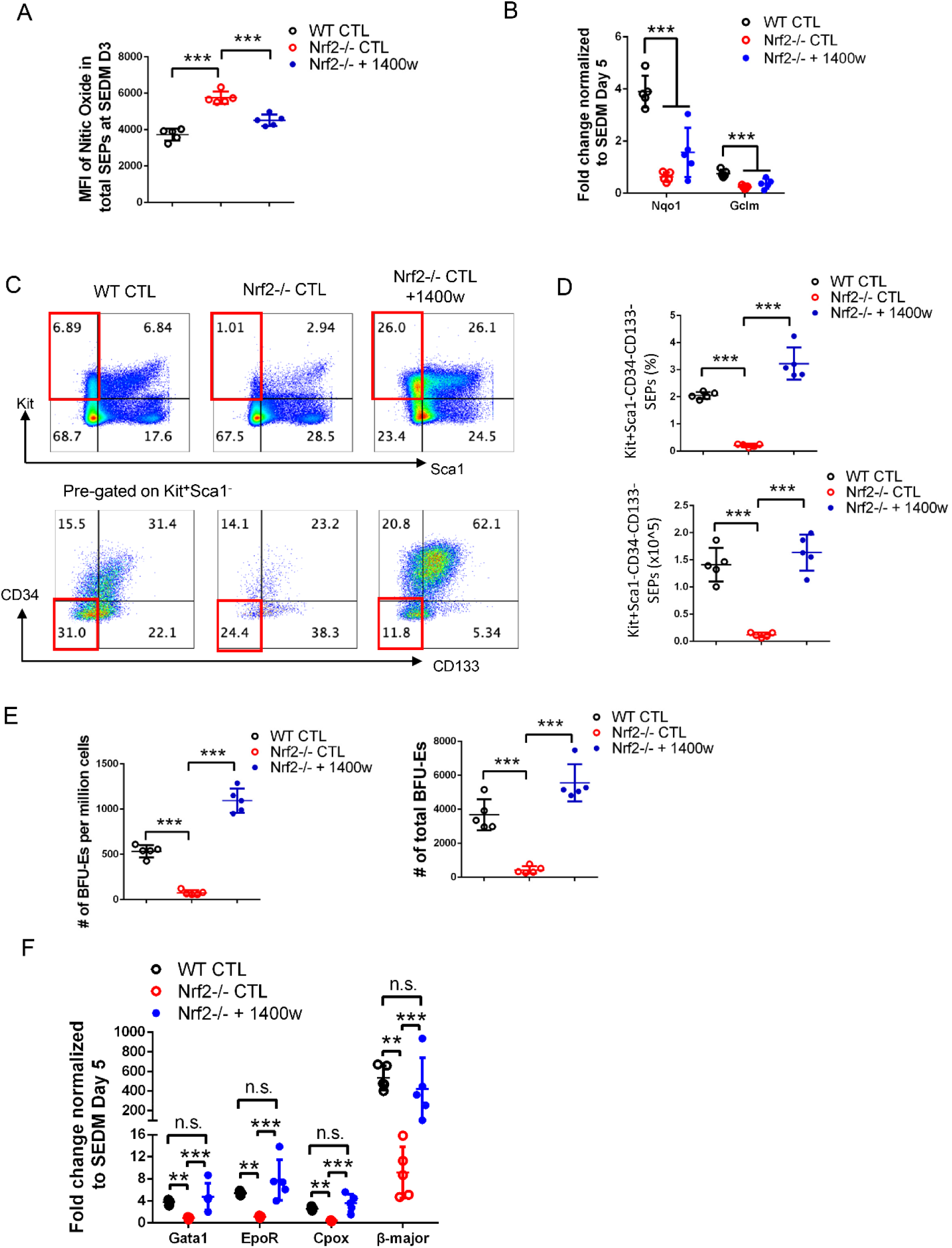
Nrf2 promotes differentiation by reducing iNOS-derived NO production. (A-F) WT and Nrf2-/-BM cells were cultured in SEEM for 5 days, after which were switched to SEDM. Nrf2-/-SEDM cultures were treated ± 10 μM 1400w for 3 days. Quantification of the intracellular NO levels by MFI of DAF-FM DA staining (A). qRT-PCR detection of mRNA abundance of Nrf2-target genes *Nqo1* and *Gclm* (B). Representative flow cytometry plot showing Kit+Sca1-CD34-CD133-cells (C), and quantification of percentages (top) and absolute numbers (bottom) of Kit^+^Sca1^-^CD34^-^CD133^-^ differentiating SEPs (D). Quantification of frequency (left) and absolute numbers (right) of stress BFU-Es by colony assay (E). qRT-PCR detection of mRNA abundance of indicated erythroid-specific genes (F) (n=5, one-way ANOVA/Tukey’s). Data represent mean ± SD. n.s. p > 0.05, ** p < 0.01, *** p < 0.001.

## DISCUSSION

Stress erythropoiesis is a component of the inflammatory response, which compensates for the loss of steady-state erythroid production to maintain homeostasis during times of infection and tissue damage. Our previous work showed that pro-inflammatory signals play a key role in the proliferation of SEPs(^9^). Here we build on our earlier data to show that induction of NO production promotes a metabolic state that drives proliferation through the activation of anabolic pathways. The result is the expansion of a transient amplifying population of SEPs. NO is produced by SEPs and niche cells through increased activity of the AASS pathway and usage of glutamine. Our previous work showed that GDF15 and Yap1-dependent signaling are required for the efficient expansion of immature SEP populations. Mutation in either of these genes significantly decreases the expression of enzymes involved in glutamine metabolism, *Gls1* and *Got1*. The defects in proliferation observed in these mutants were partially rescued by adding exogenous glutamate to the cultures(^28, 31^). Our data expand on the observations of Oburoglu et al. who showed that the proliferation of erythroid progenitors in the spleens of PHZ-treated mice was inhibited by DON treatment, which is consistent with our observation that Nos2-/-mice exhibit a defective expansion of SEPs in spleen after PHZ-induced anemia. Those studies also showed that nucleotide biosynthesis was defective in DON-treated cultures (^20, 49^).

Similarly, our data show that inhibition of iNOS with 1400w leads to a defect in nucleotide biosynthesis (Figure 2H-J). Taken together our study builds on these previous studies to show that glutamine metabolism driving NO production plays a key role in the expansion of SEPs.

NO can S-nitrosylate proteins on cysteine residues(^50^) and many enzymes in the glycolytic, TCA and oxidative phosphorylation pathways are known targets suggesting that NO dependent modifications can regulate enzyme activity(^51^). These data suggest that the effect of NO in proliferating SEPs is to alter the activity of enzymes in the glycolytic, PPP and serine/glycine pathways to support rapid cell division. Inhibition of iNOS with 1400W altered the flux of labeled carbons through the late glycolytic cycle as levels of labeled PEP increased while labeled pyruvate and lactate decreased, which is consistent with previous work showing NO regulates Pkm2 (Figure 2F)(^39, 40^). Similarly, 1400W-treatment led to increased levels of labeled 3-PG while levels of serine and glycine are reduced suggesting that NO regulates the serine/glycine pathway (Figure 2L). Future studies will be needed to understand the exact nature and consequences of NO dependent modifications on the metabolism in these cells.

The identification of NO as a key signal promoting the proliferation of SEPs has implications for human anemia. Sickle cell disease (SCD) patients have limited NO bioavailability (^52^). Treatment of SCD patients with hydroxyurea improves erythropoiesis by increasing the production of fetal hemoglobin, HbF. Hydroxyurea acts as a NO donor, nitrosylating the soluble guanylate cyclase, sGC(^53^). We previously showed that SEPs can be readily cultured from the peripheral blood of SCD patients, but not healthy controls. Despite the active mobilization of stress erythropoiesis, these patients exhibit anemia(^6^). These data suggest that low NO levels could also affect the proliferation of SEPs in patients, compromise the efficiency of stress erythropoiesis and exacerbate the inefficient steady-state erythropoiesis already observed in SCD patients(^54^).

In summary our data show that the rapid expansion of immature SEPs is driven by NO-dependent changes in metabolism that favor anabolic processes. This signaling is antagonized by the activation of Nrf2, leading to a down regulation of *Nos2* expression and a decrease in NO levels. The metabolic state induced by NO inhibits the expression of the erythroid program and the role of Nrf2 in inducing erythroid differentiation is to inhibit NO production.

## Acknowledgments

We thank the members of the Paulson lab for suggestions and comments on the work, especially Yuting Bai for her help with breeding and maintaining mice. We thank Sougat Misra for helpful discussion about metabolism, Philip Smith and Justin Munro for running LC-MS, and Erik Allman for his advice on isotope tracing study design. The visual abstract was created with BioRender.com. This work was supported by NIH grant HL146528 (RFP), NIFA-USDA Hatch Project PEN04771 accession #0000005 (RFP and KSP), R01CA239256 (MH), NIFA-USDA Hatch Project PEN04275 accession #1018544 (MH), startup funds from the College of Agricultural Sciences, Pennsylvania State University (MH), the Dr. Frances Keesler Graham Early Career Professorship from the Social Science Research Institute, Pennsylvania State University (MH), the NIFA-USDA Hatch Project PEN04607 accession number 1009993 (ADP) and from the PA Department of Health using Tobacco CURE funds (ADP, IK).

## Authorship

B.R., K.S.P and R.F.P. conceived and designed the study. B.R., S.T. and Y.C. performed the experiments. B.R., J.M., and M.A.H. analyzed the RNA-seq data. B.R., I.K., J.C., and A.D.P. analyzed the metabolomics analysis. B.R. and R.F.P. wrote the initial draft of the manuscript, and all authors were involved in review and editing.

Conflict-of-interest disclosure: The authors declare no competing financial interests.

