## Supplemental methods and data for "Nitric oxide regulates metabolism in murine stress erythroid progenitors to promote recovery during inflammatory anemia"

Ruan et al.

Supplemental Methods and Data

### Supplemental Methods

#### Murine stress erythropoiesis culture

Isolated murine bone marrow cells were cultured in stress erythropoiesis expansion media (SEEM) at a starting concentration of 6 x $10^5$ cells/ml for 5 to 7 days and switched to stress erythropoiesis differentiation media (SEDM) at 3 x $10^5$ cells/ml for 3 days.

SEEM was prepared by supplementing IMDM with 10% (*v/v*) FBS, 0.0007% (*v/v*) 2-mercaptoethanol, 0.01 g/ml BSA,10 µg/ml ciprofloxacin, 2 mM L-glutamine, 10 µg/ml insulin (Sigma-Aldrich), 200 µg/ml holo-transferrin (Alfa Aesar), 50 ng/ml murine SCF (GoldBio), 15 ng/ml human BMP4 (Thermo Fisher Scientific), 25 ng/ml murine SHH (GoldBio) and 30 ng/ml murine GDF15 (Biomatik). SEDM was prepared by additionally supplementing SEEM with 3 U/ml human Epo. SEEM were cultured in ambient air (20% $O_2$), while SEDM were cultured in a hypoxia chamber (2% $O_2$) unless otherwise noted. All cultures were incubated in 5% $CO_2$ at 37 °C.

#### Human stress erythropoiesis culture

Primary human bone marrow mononuclear cells (BMNCs) purchased from AllCells were used for human stress erythropoiesis cultures. BMNCs were adjusted to a concentration of 1 x $10^6$ cells/ml in human SEEM and cultured in 37°C with 5% $CO_2$ and 20% $O_2$ for 7 days. Human SEEM was prepared by supplementing IMDM with 10% (*v/v*) embryonic stem-cell FBS, 0.0007% (*v/v*) 2-mercaptoethanol, 20% (*v/v*), BIT 9500 serum substitute (STEMCELL Technologies), 10 µg/ml ciprofloxacin, 2 mM L-glutamine, and 50 ng/ml human SCF (GoldBio), 15 ng/ml human BMP4 (Thermo Fisher Scientific), 25 ng/ml human SHH (GoldBio) and 30 ng/ml human GDF15 (R&D Systems).

#### Phenylhydrazine (PHZ)-induced hemolytic anemia

To induce acute hemolytic anemia, age- and sex-matched mice were injected intraperitoneally with a single dose (100 mg/kg body weight) of freshly prepared phenylhydrazine (Sigma-Aldrich, dissolved in PBS). Stress erythropoiesis was established to recover from anemia in the following 7 days, and mice were monitored daily for health and survival. To assess stress erythropoiesis, mice were sacrificed at indicated time points for blood (cardiac puncture) and spleen sample collection.

**_Brucella abortus_-induced inflammatory anemia**

Heat-killed *Brucella abortus* (HKBA, strain 1119-3) was used to induce anemia of inflammation following a previously described method[(1)]. To induce stress erythropoiesis, age- and sex-matched mice were administered with $5 \times 10^8$ particles of HKBA via intraperitoneal injection. In the following 28 days, mice were monitored daily for survival and health, and blood was collected retro-orbitally in every other day for microhematocrit test.

**Bone marrow transplant**

Prior to irradiation, WT and Nrf2-/- recipient mice were fed with acidified water (PH=3.0) for 1 week, followed by antibiotic water treatment for 3 days. Recipient mice were lethally irradiated at a single dose of 950 cGy, and then transplanted with $5 \times 10^5$ donor bone marrow cells in 150 µl PBS by retro-orbital injection. Recipient mice were transplanted with donor cells isolated from the same strain (WT to WT; Nrf2-/- to Nrf2-/-). After the transplantation, mice were monitored daily for survival and health, and the peripheral blood was collected retro-orbitally in every three days for hematocrit measurement.

**Complete blood count test**

Mouse blood was collected retro-orbitally with a $K_2$EDTA-coated Microtainer tube, and sample was immediately analyzed on a Hemavet 950 analyzer (Drew Scientific).

**Stress BFU-E colony assay**

$2.5 \times 10^5$ cells of isolated splenocytes or SEPs from SEDM cultures were resuspended in 2 ml MethoCult M3334 media (STEMCELL Technologies) supplemented additionally with 50 ng/ml SCF (GoldBio) and 15 ng/ml BMP4 (Thermo Fisher Scientific). Cell suspension was evenly seeded into a 12-well plate with technical triplicates. After a 5-day culture in 37 °C with 2% $O_2$ and 5% $CO_2$, cells were stained with benzidine to quantify stress BFU-E colonies.

**NO staining**

Intracellular NO levels were quantified by flow cytometry analysis using the fluorescent probe DAF-FM diacetate (Thermo Fisher Scientific) following manufacturer’s protocol. To assess NO levels in different SEP populations, cells were additionally stained for progenitor cell surface markers as described below.

**Flow cytometry analysis**

Splenocytes or non-adherent cells from SEEM or SEDM cultures were harvested. Zombie Yellow Fixable Viability Kit (BioLegend) was used to exclude non-viable cells. For analysis of mouse SEPs, cells were stained with fluorophore-conjugated cell surface antibodies against Kit, Sca1, CD34 and CD133. For analysis of human SEPs, cell surface antibodies against Kit, CD34 and CD133 were used. Flow cytometry analysis was performed on a BD LSR Fortessa Cytometer (BD Biosciences) and data were analyzed by FlowJo software (BD Biosciences). See Supplemental Table 1 for flow cytometry antibodies.

**Metabolite extraction**

Cell pellets were extracted with 1 ml pre-chilled 50:50 HPLC-grade water:methanol (*v/v*) containing 1 μM chlorpropamide as an internal standard. The samples were homogenized and were then snap frozen with liquid nitrogen and immediately thawed at room temperature. This step was repeated for three times followed by centrifuging for 10 min at 12,000 x g and 4 °C. The supernatants were transferred into fresh microfuge tubes. The remaining cell pellets were re-extracted with 0.5 ml 50% methanol containing 1 μM chlorpropamide as described above. The supernatants were then combined with the first extraction. Metabolites-containing supernatants were concentrated to dryness at room temperature in a SpeedVac concentrator and re-dissolved in 100 μl 97:3 water:methanol (*v/v*). After centrifuging for 10 min at 13000 × g and 4°C, 70 μl of supernatants were transferred into autosampler vials for LC-MS analysis. Two types of control were prepared in triplicates to run in concert with the experimental samples: the process blank control, and the pooled control containing an equal volume from each experimental sample.

**LC-MS analysis**

The sample run order was randomized to reduce bias from instrument drift. 10 µl sample was subjected to LC-MS analysis on a Exactive Plus Orbitrap mass spectrometer (Thermo Fisher Scientific) coupled to an Ultimate 3000 UHPLC system (Thermo Fisher Scientific). Reversed-phase chromatography mode was used to separate compounds on a Xselect C18 HSS column (Waters) with solvent A (97:3 water:methanol (*v/v*), 10 mM tributylamine, and 15 mM acetic acid ) and solvent B (methanol). The flow rate was 200 µl/min, and the total run time was 25 min. The gradient was 0 min, 0% B; 5 min, 20% B; 7.5 min, 55% B; 15 min, 65% B; 17.5 min, 95% B; and 21 min, 0% B. The mass spectrometer was operated using electrospray ionization method in a negative-ion mode and was at a resolution of 140,000 FWHM with a scan range of 85 to 1000 m/z.

**$^{13}C$ isotope tracing and metabolic profiling**

Untreated SEEM cultures at day 1,3 and 5, and 1400w-/DMSO-treated SEEM cultures (10 µM at day 3 for 48 hrs) were re-cultured in SEEM supplemented with 25 mM U-[$^{13}C$]-Glucose for another 24 hrs. Extracted metabolites were resuspended in 3% methanol (*v/v*). Samples were subjected to LC-MS analysis on a Exactive Plus Orbitrap mass spectrometer coupled to an Ultimate 3000 UHPLC system (Thermo Fisher Scientific) as described previously[(2)]. Raw data files when converted to .mzML format[(3)] were analyzed by MS-DIAL software[(4)]. Metabolites were identified by comparison to an in-house reference library of pure metabolites. Peak areas of identified metabolites were normalized to internal standard and cell numbers.

**RNA sequencing**

Samples prepared in biological triplicates were submitted to Beijing Genomics Institute (BGI Group) for RNA sequencing. Filtered clean reads were aligned to the GRCm38 mouse reference genome with HISAT2[(5)]. The output was processed by SAMtools to generate sorted

BAM files. Gene expression was quantified by StringTie[6]. The output was processed according to the StringTie manual before import into DESeq2 for differential gene expression analysis[7]. Gene annotation and enrichment analysis were based on GO and KEGG database and were performed using ClusterProfiler[8].

**qRT-PCR**

Total RNA was extracted from spleen cells or cultured progenitors with TRIzol reagent (Thermo Fisher Scientific) according to manufacturer's guidance. RNA concentration was quantified using a NanoDrop One Spectrometer (Thermo Fisher Scientific), and 1 µg of total RNA was subjected to cDNA preparation using the qScript cDNA Synthesis Kit (Quanta Biosciences). TaqMan Gene Expression assays were used for qRT-PCR analysis. qPCR was performed on a StepOnePlus Real-Time PCR System (Applied Biosystems) using the PerfeCTa qPCR SuperMix ROX (Quanta Biosciences). Gene expression was quantified by the $\Delta\Delta C_T$ method in reference to the housekeeping gene 18S rRNA for normalization. See Supplemental Table 3 for TaqMan probes.

**Isolation SEPs from the spleen for metabolomics analysis.**

Murine spleens were harvested and disassociated to single cell suspension at a concentration of 1 x 10^8 cells/mL. Labeling reagent was added at a concentration of 50 µL/mL of sample. Cells were mixed and incubated at RT for 15 mins. Selection cocktail was added to sample at a 70 µL/mL of sample. Cells were mixed and incubated at RT for 15 mins. RapidSpheres were added to sample at a 50 µL/mL of sample. Cells were mixed and incubated at RT for 10 mins. Cells were washed with FACS buffer for 4 times and were resuspended for downstream metabolomics analysis.

**Supplemental Table 1. Flow cytometry antibody list**

| Antibodies | Source | Identifier |
|---|---|---|
| Brilliant Violet 421 anti-mouse CD117 (c-Kit), Clone 2B8 | BioLegend | Cat# 105828; RRID: AB_11204256 |
| APC/Cyanine7 anti-mouse Ly-6A/E (Sca-1), Clone D7 | BioLegend | Cat# 108126; RRID: AB_10645327 |
| FITC anti-mouse Ly-6A/E (Sca-1), Clone D7 | BioLegend | Cat# 108106; RRID: AB_313343 |
| PE/Cyanine7 anti-mouse CD133, Clone 315-2C11 | BioLegend | Cat# 141210; RRID: AB_2564069 |
| Alexa Fluor 647 anti-mouse CD34, Clone RAM34 | BD Biosciences | Cat# 560230; RRID: AB_1645200 |
| FITC anti-mouse CD34, Clone RAM34 | BD Biosciences | Cat# 553733; RRID: AB_395017 |
| FITC anti-mouse CD45.2, Clone 104 | BD Biosciences | Cat# 553772; RRID: AB_395041 |
| Brilliant Violet 421 anti-human CD117 (c-kit), Clone 104D2 | BioLegend | Cat# 313216; RRID: AB_11148721 |
| Alexa Fluor 647 anti-human CD34, Clone 561 | BioLegend | Cat# 34361; RRID: AB_2632632 |
| PE anti-human CD133, Clone 7 | BioLegend | Cat# 372804; RRID: AB_2632880 |

**Supplemental Table 2. Software and algorithm list**

| | |
|---|---|
| GraphPad Prism version 6.07 | https://www.graphpad.com |
| R version 4.0.3 | https://www.r-project.org |
| Cytoscape version 3.8.0 | https://cytoscape.org |
| FlowJo version 10.8.0 | https://www.flowjo.com |
| ImageJ version 1.52 | https://imagej.nih.gov/ij/index.html |
| ProteoWizard version 3 | https://proteowizard.sourceforge.io |
| MS-DIAL version 3.40 | http://prime.psc.riken.jp/compms/msdial/main.html |
| HISAT2 version 2.2.0 | http://daehwankimlab.github.io/hisat2/ |
| SAMtools version 1.4 | http://www.htslib.org |
| StringTie version 2.1.4 | https://ccb.jhu.edu/software/stringtie/ |
| DESeq2 version 1.30.1 | https://bioconductor.org/packages/release/bioc/html/DESeq2.html |
| ClusterProfiler version 3.18.0 | https://bioconductor.org/packages/release/bioc/html/clusterProfiler.html |

**Supplemental Table 3. List of TaqMan probes for qRT-PCR.**

| Probes and primers | Identifier |
|---|---|
| Human 18S | Hs99999901_s1 |
| Mouse Nos2 | Mm00440502_m1 |
| Mouse Gls | Mm01257297_m1 |
| Mouse Got1 | Mm01195792_g1 |
| Mouse Asl | Mm01197741_m1 |
| Mouse Ass1 | Mm00711256_m1 |
| Mouse Arg1 | Mm00475988_m1 |
| Mouse Nfe2l2 | Mm00477784_m1 |
| Mouse Nqo1 | Mm01253561_m1 |
| Mouse Gata1 | Mm01352636_m1 |
| Mouse EpoR | Mm00833882_m1 |
| Mouse Cpox | Mm00483982_m1 |
| Mouse Gclm | Mm01324400_m1 |
| Mouse β Major Probe | 6FAM- CTCTCTTGGGAACAATTAACCATTGTTCACAG-TAMRA |
| Mouse β Major Forward Primer | 5’ -AACCCCCTTTCCTGCTCTTG- 3’ |
| Mouse β Major Reverse Primer | 5’ -TCATTTTGCCAACAACTGACAGA- 3’ |
| Mouse β H1 Probe | 6FAM- ACTTTCTTGCCATGGGCTCTAATCCGG-TAMRA |
| Mouse β H1 Forward Primer | 5’ -CCTGGCCATCATGGGAAAC- 3’ |
| Mouse β H1 Reverse Primer | 5’-CCCCAAGCCCAAGGATGT-3’ |

**Supplemental method reference**

### Supplemental Figure 1

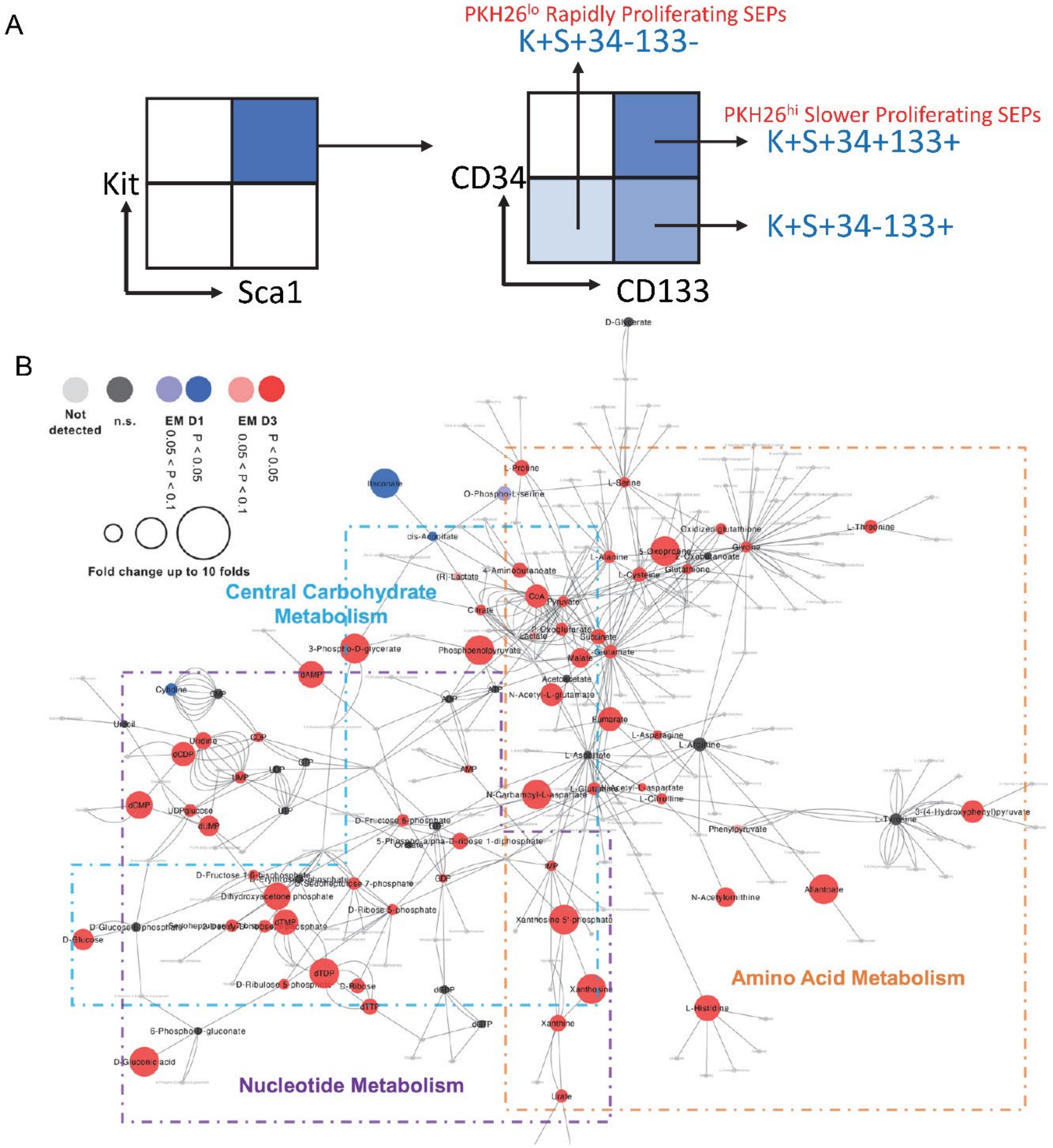

C

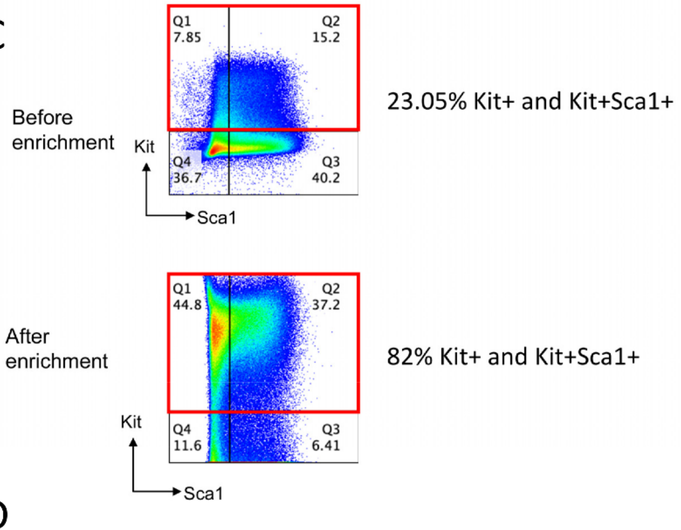
Before
enrichment
Q1
7.85
Q2
15.2
Q4
36.7
Q3
40.2
Kit
Sca1
23.05% Kit+ and Kit+Sca1+
After
enrichment
Q1
44.8
Q2
37.2
Q4
11.6
Q3
6.41
Kit
Sca1
82% Kit+ and Kit+Sca1+

D

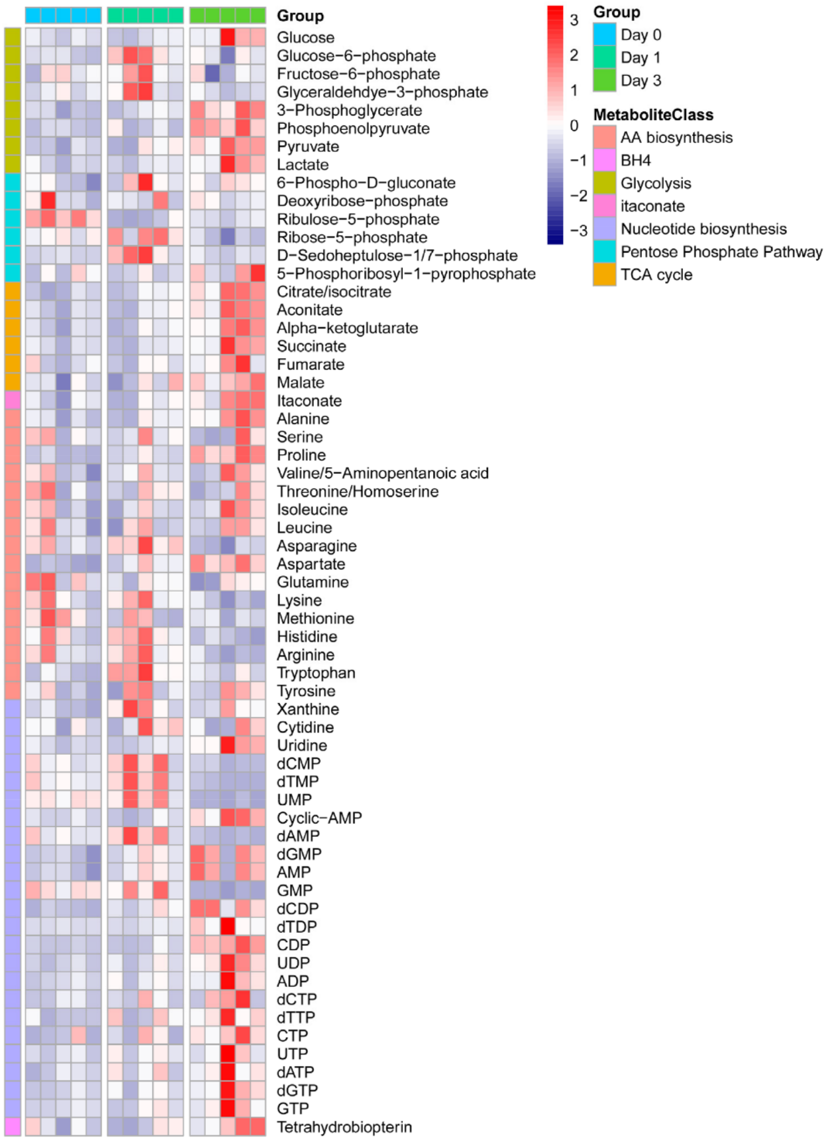
Group
Glucose
Glucose−6−phosphate
Fructose−6−phosphate
Glyceraldehdye−3−phosphate
3−Phosphoglycerate
Phosphoenolpyruvate
Pyruvate
Lactate
6−Phospho−D−gluconate
Deoxyribose−phosphate
Ribulose−5−phosphate
Ribose−5−phosphate
D−Sedoheptulose−1/7−phosphate
5−Phosphoribosyl−1−pyrophosphate
Citrate/isocitrate
Aconitate
Alpha−ketoglutarate
Succinate
Fumarate
Malate
Itaconate
Alanine
Serine
Proline
Valine/5−Aminopentanoic acid
Threonine/Homoserine
Isoleucine
Leucine
Asparagine
Aspartate
Glutamine
Lysine
Methionine
Histidine
Arginine
Tryptophan
Tyrosine
Xanthine
Cytidine
Uridine
dCMP
dTMP
UMP
Cyclic−AMP
dAMP
dGMP
AMP
GMP
dCDP
dTDP
CDP
UDP
ADP
dCTP
dTTP
CTP
UTP
dATP
dGTP
GTP
Tetrahydrobiopterin
3
2
1
0
−1
−2
−3
Group
Day 0
Day 1
Day 3
MetaboliteClass
AA biosynthesis
BH4
Glycolysis
itaconate
Nucleotide biosynthesis
Pentose Phosphate Pathway
TCA cycle

**Supplemental Figure 1. Metabolism is rewired to increase anabolic synthesis for SEP proliferation.**

(A) Gating strategy for flow cytometry analysis of proliferating SEPs. Cells were stained for viability followed by gating on Kit and Sca1. Pre-gated Kit+Sca1+ cells were then gated on CD34 and CD133.

(B) Cytoscape network analysis comparing metabolites between SEEM day 1 and day 3 SEPs in central carbohydrate, nucleotide and amino acid metabolism. Red indicates metabolite level higher in day3, blue indicates higher in day 1 and grey indicates no difference or not detected. Circle size represents fold change (n=5 per time point).

(C) Mice were treated with HKBA, and Kit+ cells were isolated from splenocytes at day 8 using the EasySep mouse CD117 positive selection kit. Flow cytometry analysis of SEPs before and after Kit enrichment.

(D). Mice were treated with PHZ, on the indicated days after treatment, spleen cells were isolated. Kit+ cells were isolated using the EasySep mouse CD117 positive selection kit (Stem Cell technologies, Vancouver, BC). SEPs were processed for metabolomics analysis as described in the methods. (Top) a representative flow diagram showing that Kit+ cells isolated using the Kit contained all the SEPs in the spleen. (Bottom) Heat map showing metabolites levels. N=5 for each day.

### Supplemental Figure 2

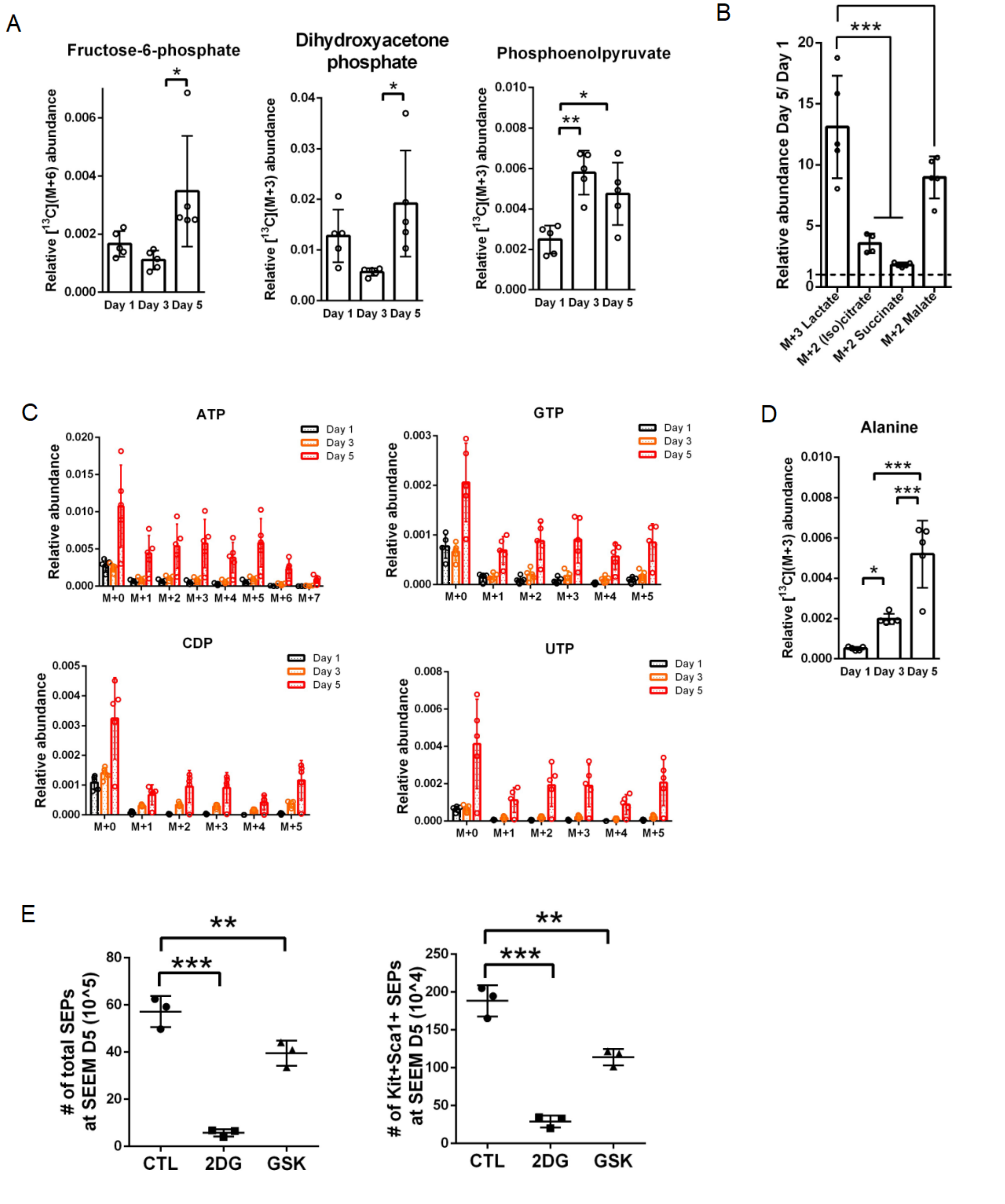

**Supplemental Figure 2. $^{13}C$-isotope tracing analysis of progenitors in SEEM cultures over time.**

(A-D) SEPs isolated from regular SEEM on day 1, 3 and 5 were re-cultured in SEEM supplemented with 25 mM U-[$^{13}C$]-Glucose. After another 24 hrs, SEPs were collected for metabolic profiling. Abundance of $^{13}C$-isotope labeling of selected metabolites in glycolysis on day 5 relative to day1 (A). Isotope tracing of lactate and TCA cycle intermediates (B), nucleotide metabolism (C) and alanine (D) (n=5 per time point, one-way ANOVA/Holm-Sidak).

(E) SEEM cultures were treated with vehicle, 1 mM 2DG or 13 nM GSK at day 3 for 48 hrs. Total SEP cell counts (left), and flow cytometry quantification of absolute numbers of $Kit^{+}Sca1^{+}$ SEPs (right) (n=3 per group, one-way ANOVA/Dunnett's).

Data represent mean ± SEM. * $p < 0.05$, ** $p < 0.01$, *** $p < 0.001$.

**Supplemental Figure 3**

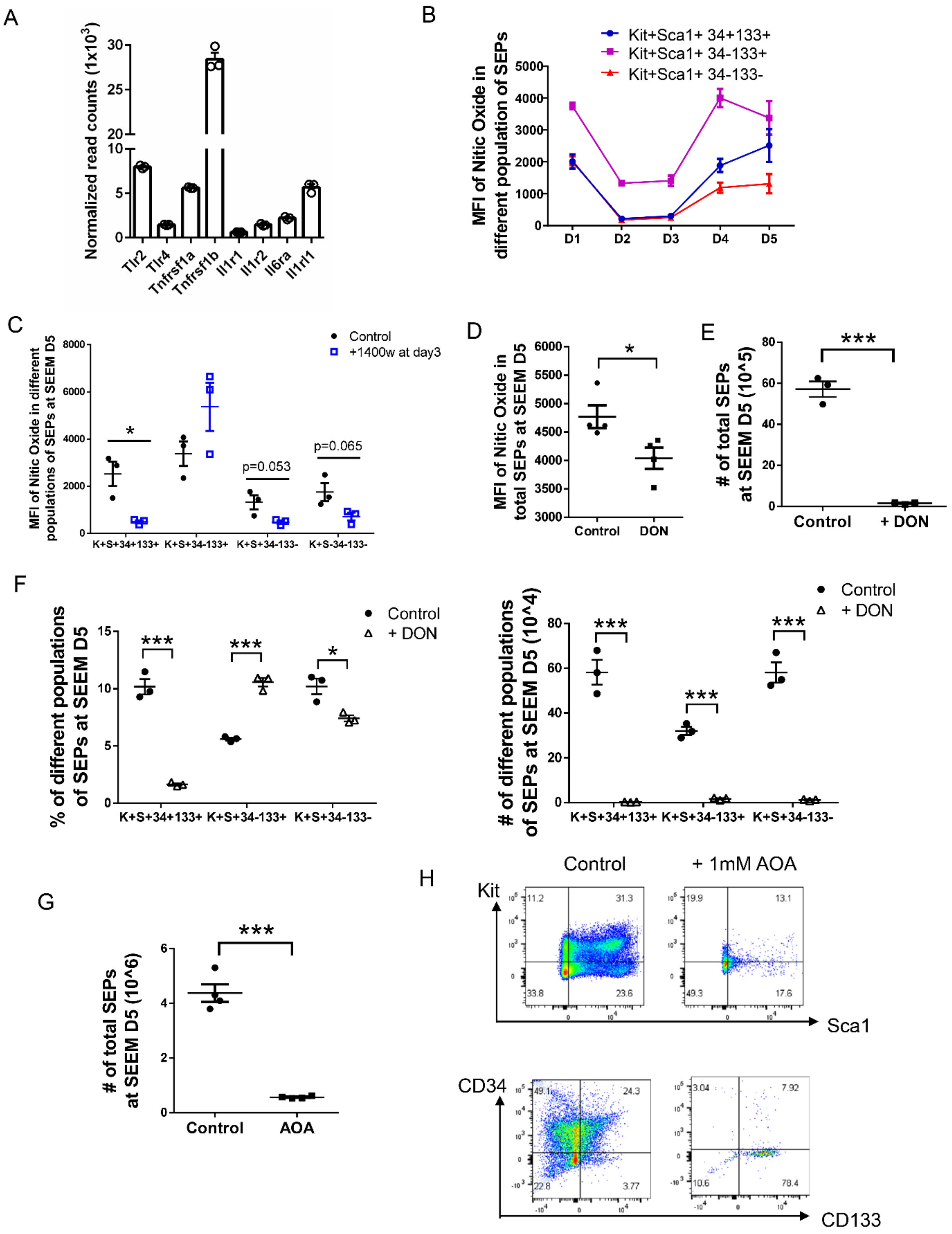

**Supplemental Figure 3. Glutamine metabolism contributes to increased AASS flux to fuel NO production.**

(A) RNA-seq data showing mRNA abundance of TLRs and receptors for pro-inflammatory cytokines in SEPs isolated at SEEM day 5 (n=5).

(B) NO levels of indicated SEP populations in SEEM cultures over time quantified by MFI of DAF-FM DA staining (n=3 per time point).

(C) Quantification of NO levels in indicated SEPs grown in SEEM cultures treated ± 10 μM 1400w for 48 hrs (n=3 per group, unpaired t test).

(D) SEEM cultures were treated treated ± 20 μM DON at day 5 for 3 hrs, followed by quantification of NO levels (n=4 per group, unpaired t test).

(E-F) SEEM cultures were treated ± 20 μM DON at day 3 for 48 hrs. Quantification of total numbers of SEPs (E), and flow cytometry analysis of the percentages (left) and absolute numbers (right) of indicated populations of SEPs (F) (n=3 per group, unpaired t test; shared same negative control with supplemental Figure 1C and Figure 1K).

(G-H) SEEM cultures were treated ± 1 mM AOA at day 3 for 48 hrs. Quantification of total numbers of SEPs (G) and flow cytometry analysis of different populations of SEPs (H) (n=4, unpaired t test).

Data represent mean ± SEM. * $p < 0.05$, *** $p < 0.001$.

### Supplemental Figure 4

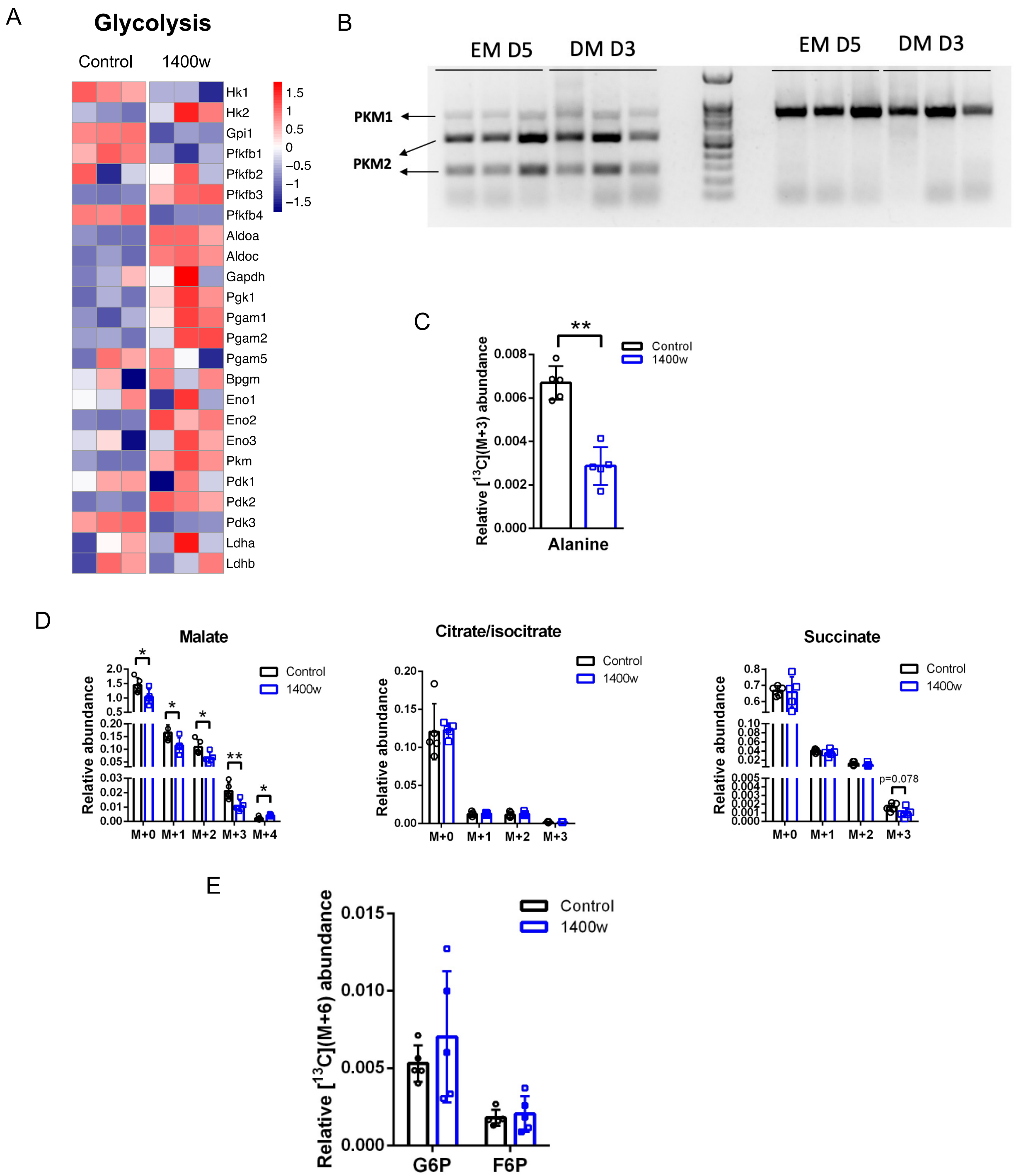

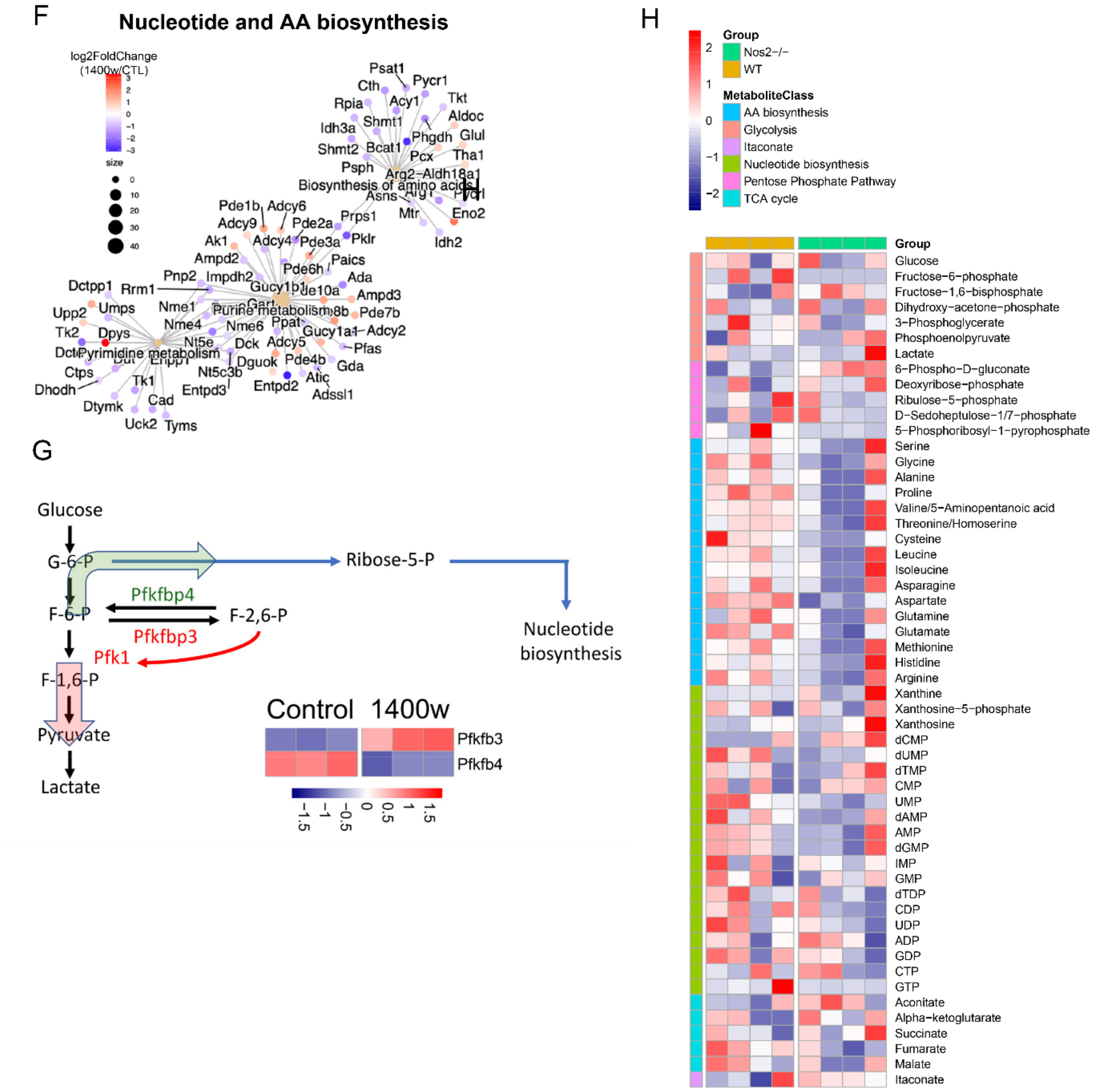


**Supplemental Figure 4. Pro-inflammatory iNOS signaling contributes to glycolysis and anabolic synthesis at the expansion stage of stress erythropoiesis.**

(A) Heat map from RNA-seq analysis comparing SEPs treated ± 10 μM 1400w at SEEM day 3 for 48 hrs. Analysis depicting the mRNA abundance of glycolytic enzymes. Color bar represents row-wise scaled z-score (n=3 per group). (B) Pkm2 is the dominant Pkm isoform in SEPs. The alternative exons were amplified from mRNA by RT-PCR. (left) PCR products were cut with Pst1, which cuts the Pkm2 sequence. (right) uncut control. EM D5 is SEEM culture at day 5, DM

D3 is SEDM Culture on day 3. N=3 for SEEM and SEDM. (C-E) SEPs were treated with vehicle or 10 μM 1400w at SEEM day 3 for 48 hrs, followed by [U-$^{13}$C] glucose labeling. Relative abundance of isotope labeling of alanine (C), TCA cycle intermediates (D) and selected glycolytic intermediates (E) (n=5 per group, unpaired t test). (F) Network analysis of DEGs (FDR < 0.05, |FC| > 1.2) in nucleotide and AA biosynthesis pathways with color representing log2FC of SEEM 1400w/CTL (n=3 per group). (G) Diagram of the different roles of Pfkfbp3 and Pfkfbp4 in regulating metabolite flux. RNA seq data is taken from panel A. (H). In vivo metabolomics analysis of control and Nos2-/- SEPs isolated from the spleen on day 3 after PHZ treatment. SEPs were isolated using the Kit+ EasySEP kit and isolated SEPs were processed for metabolomics analysis as described above. N=4 for each genotype.

Data represent mean ± SD. * $p < 0.05$, ** $p < 0.01$.

### Supplemental Figure 5

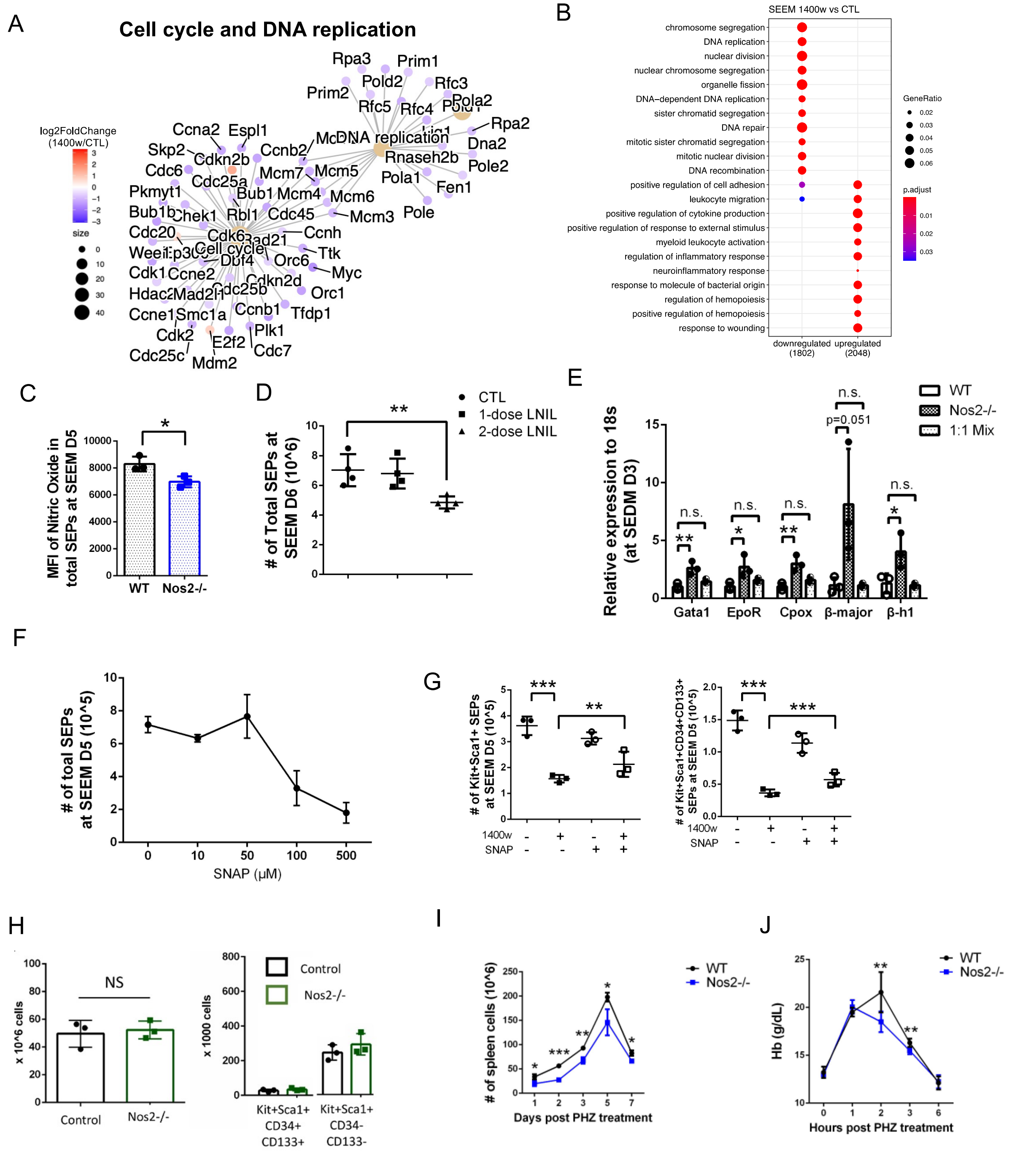


268

269

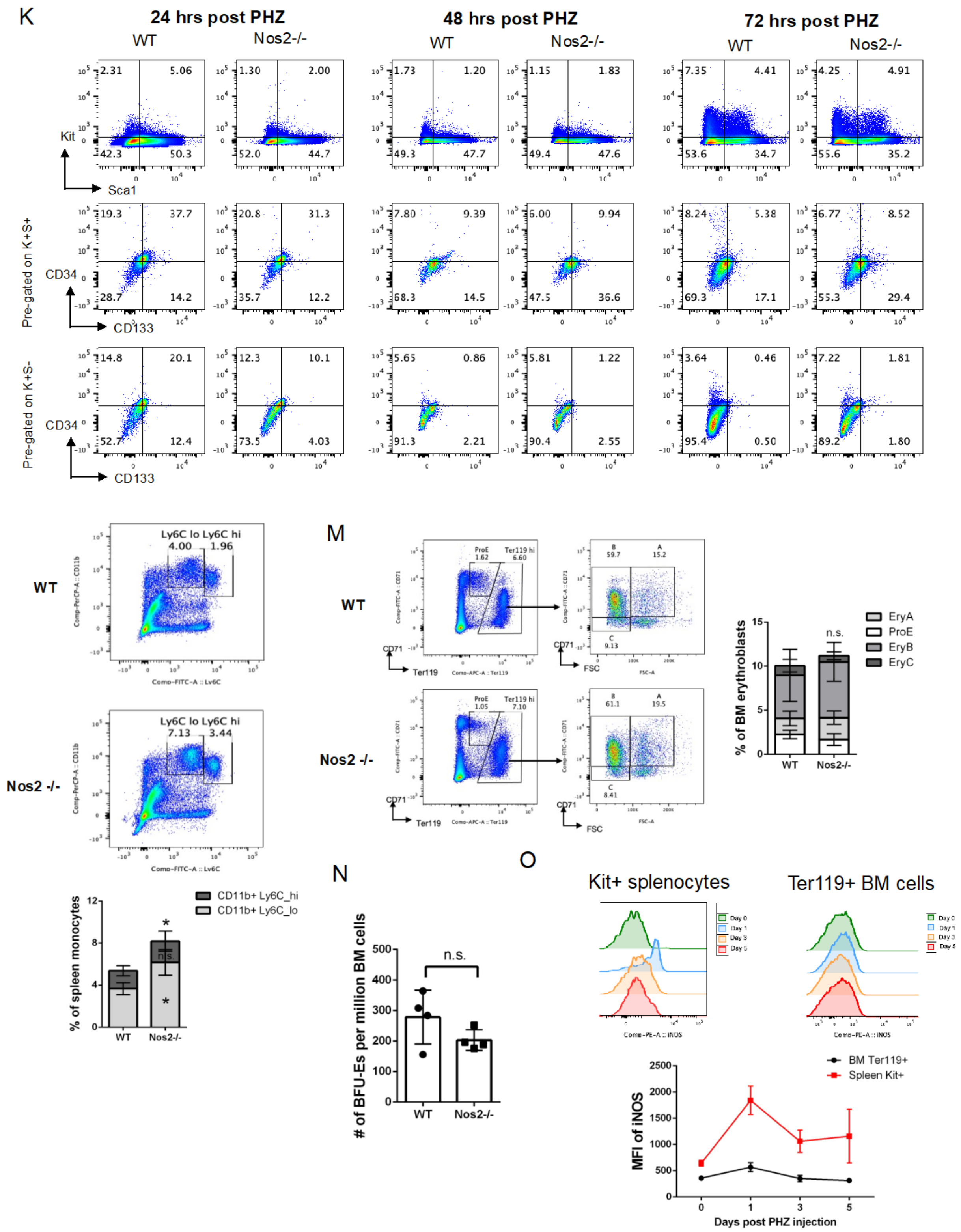


**Supplemental Figure 5. iNOS-dependent production of NO promotes SEP proliferation.**

(A-B) RNA-seq analysis comparing SEPs treated ± 10 µM 1400w at SEEM day 3 for 48 hrs.

Network analysis of DEGs (FDR < 0.05, |FC| > 1.2) in cell cycle-related pathways with color representing log2FC of SEEM 1400w/CTL (A), and top enriched GO terms identified via overrepresentation analysis of downregulated DEGs (FDR < 0.05, FC of 1400w/CTL < -1.5) and upregulated DEGs (FDR<0.05, FC > 1.5). Circle size represents gene ratio and color represents adjusted p value (B) (n=3 per group).

(C) Quantification of NO levels in SEPs grown in WT and Nos2-/- SEEM cultures (n=3 per group, unpaired t test).

(D) SEEM cultures were treated with vehicle, 1-dose (20 µg/ml) L-NIL at day 3, or 2-dose L-NIL (1st dose 10 µg/ml at day 0 and 2nd dose 20 µg/ml at day 3). Quantification of total SEP cell counts at SEEM day 6 (n=4 per group, one-way ANOVA/Dunnett's).

(E) qRT-PCR analysis of indicated erythroid-specific genes in SEPs from WT, Nos2-/- and WT/Nos2-/- 1:1 Mixed SEDM cultures (n=3 per group, one-way ANOVA/Tukey's).

(F) Total SEP cell counts in SEEM cultures treated with SNAP for 48 hrs (n=3 per group).

(G) Flow cytometry quantification of total numbers of Kit+Sca1+ (left) and Kit+Sca1+CD34+CD133+ SEPs (right) in SEEM cultures treated with 10 µM 1400w or 10 µM SNAP alone, or in combination for 24 hrs (n=3 per group, one-way ANOVA/Tukey's).

(H) Analysis of total spleen cell numbers and SEP populations (CD34+CD133+Kit+Sca1+ and CD34-CD133-Kit+Sca1+) in untreated control and Nos2-/- mice (n=3).

(I-K) Analysis of recovery from PHZ-induced hemolytic anemia. Numbers of splenocytes (I) and Hb levels (J) and representative flow cytometry plot of SEPs (K) of WT and Nos2-/- mice at indicated time points post PHZ injection (n=3 (J-K); n=7-8 (I), unpaired t test).

(L-N) Mice were injected with PHZ. Splenocytes were collected at day 3 after PHZ treatment for flow cytometry analysis of CD11b+Ly6C$^{hi}$ and CD11b+Ly6C$^{lo}$ monocyte populations (L). Bone marrow cells were collected at day 3 for flow cytometry analysis of normal erythropoiesis (M) and BFU-Es (N) (n=3).

(O) Cells were harvested at indicated time points post PHZ treatment, followed by flow cytometric quantification of Nos2 protein levels in Kit+ splenocytes and Ter119+ BM cells (n=3 per time point).

Data represent mean ± SEM. * $p < 0.05$, ** $p < 0.01$, *** $p < 0.001$.

### Supplemental Figure 6

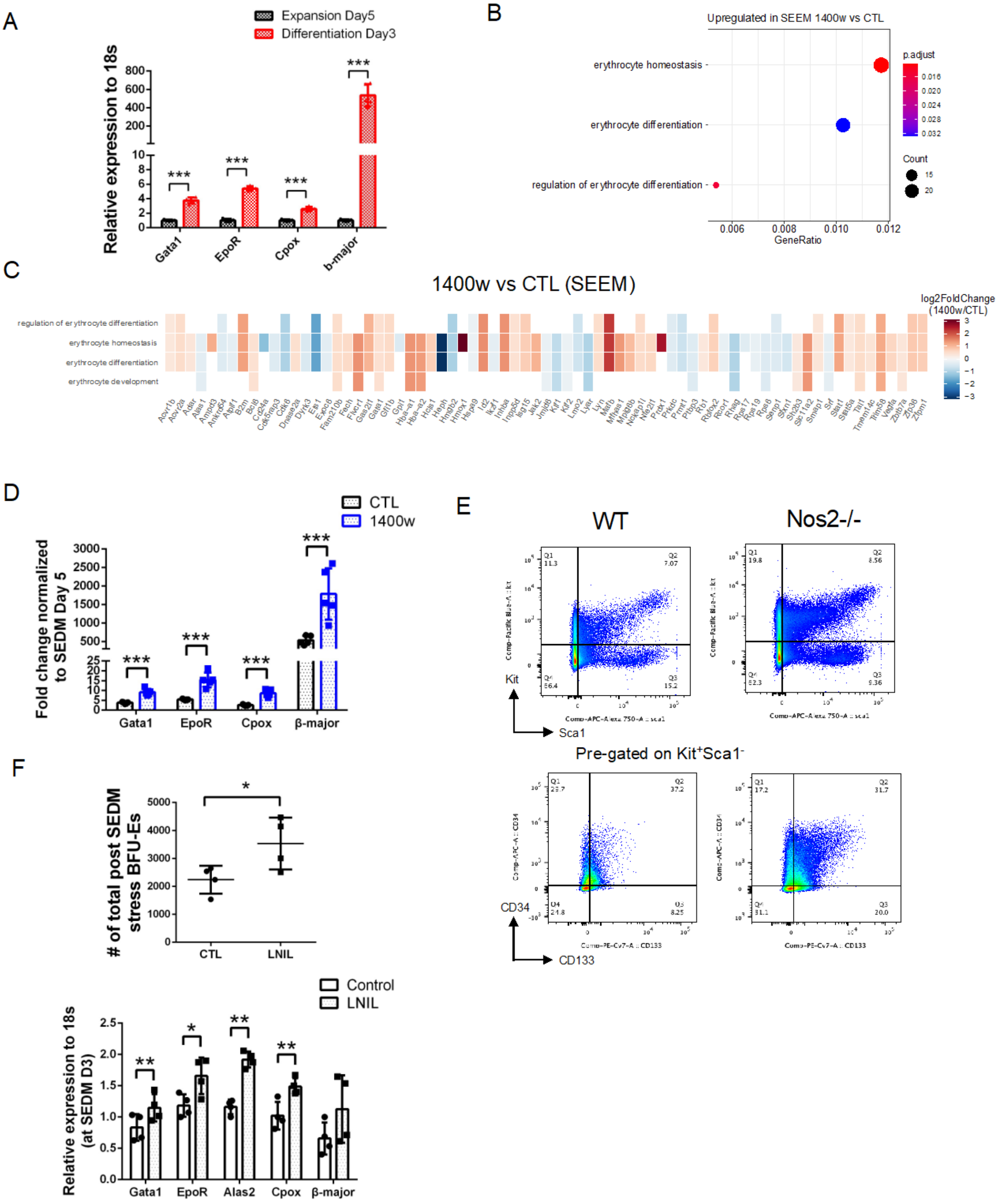

**Supplemental Figure 6. iNOS-derived NO inhibits erythroid gene expression program.**

(A) qRT-PCR analysis of indicated erythroid-specific genes between SEEM Day 5 and SEDM Day 3 (n=5 per time point, unpaired t test).

(B-C) RNA-seq analysis comparing SEPs treated with vehicle or 10 μM 1400w at SEEM day 3 for 48 hrs. Overrepresentation analysis of upregulated DEGs in SEEM 1400w (FDR < 0.05, FC of 1400w/CTL > 1.5) showing significant enriched erythroid-associated GO terms. Circle size represents the numbers of genes in each GO term and color represents adjusted p value (B). A heatmap showing DEGs (FDR < 0.05, |FC| > 1.2) in erythroid-associated pathways with color representing log2FC of SEEM 1400w/CTL (C) (n=3 per group).

(D) qRT-PCR analysis of indicated erythroid-specific genes (bottom), (n=5 per group, unpaired t test).

(E) Representative plot showing flow cytometry analysis of SEPs isolated from WT and Nos2-/- SEDM cultures at day 3.

(F) SEPs were treated with vehicle or L-NIL (20 μg/ml) when switched to SEDM cultures, followed by quantification of total stress BFU-Es by colony assay (top), and qRT-PCR analysis of indicated erythroid-specific genes (bottom) (n=4 per group, unpaired t test).

Data represent mean ± SEM. * $p < 0.05$, ** $p < 0.01$, *** $p < 0.001$.

**Supplemental Figure 7**

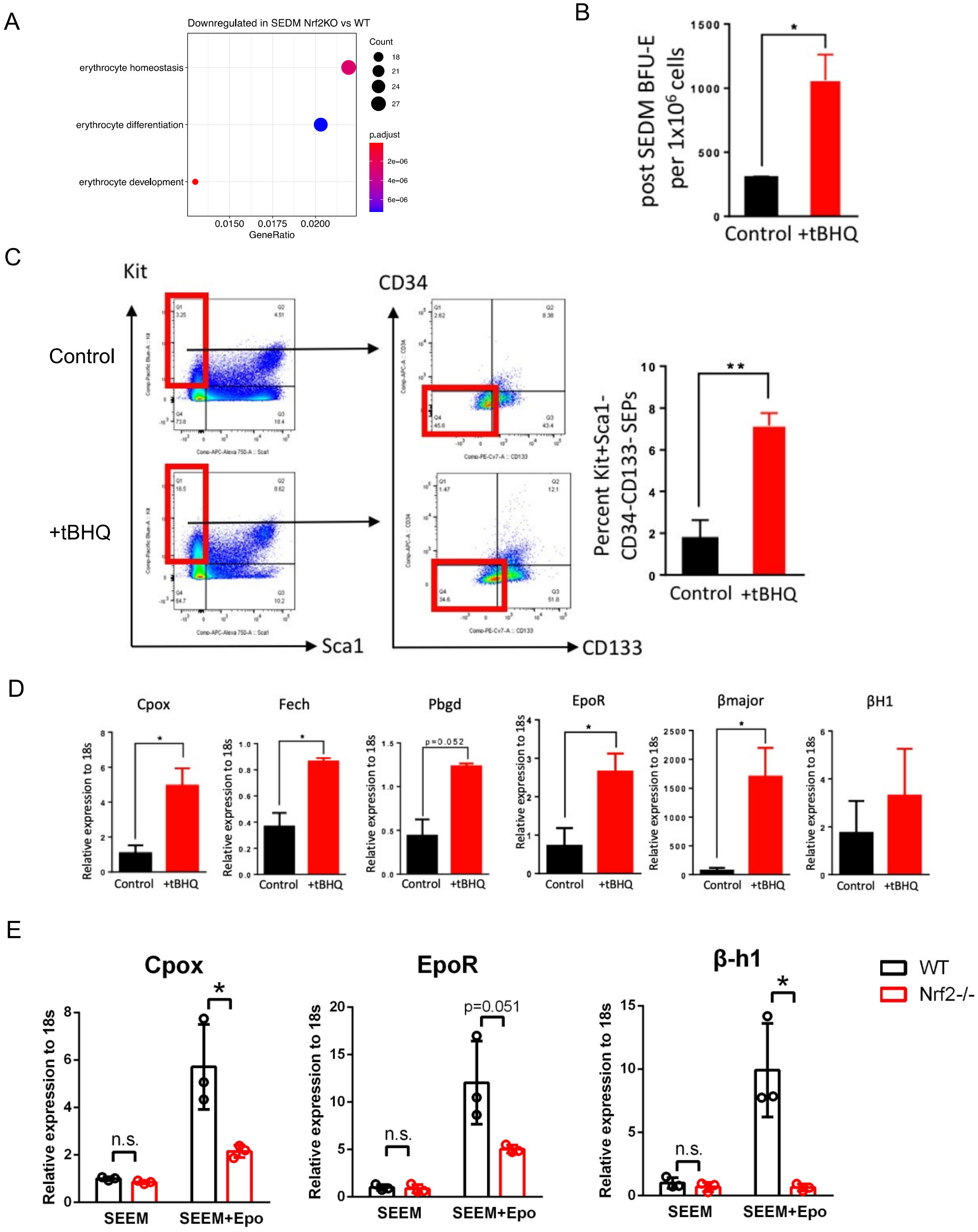

**Supplemental Figure 7. The activation of Nrf2 promotes the transition of SEPs to erythroid differentiation.**

(A) RNA-seq analysis comparing WT and Nrf2-/- SEPs isolated from SEDM cultures. Overrepresentation analysis of downregulated DEGs in SEDM Nrf2-/- (FDR < 0.05, FC of Nrf2-/-/WT < -1.5) showing erythroid-associated GO terms. Circle size represents the numbers of genes in each GO term and color represents adjusted p value (n=3 per group).

(B-D) After a 5-day culture in SEEM, SEPs were switched to SEDM and were treated with vehicle or 20μM tBHQ for 3 days. Quantification of stress BFU-Es per million cells by colony assay (B). Representative flow cytometry plot (left) and quantification of percentages of $Kit^{+}Sca1^{-}CD34^{-}CD133^{-}$ differentiating SEPs (right) (C). qRT-PCR analysis of indicated erythroid-specific genes (n=2 per group, unpaired t test).

(E) After a 5-day culture in SEEM, WT and Nrf2-/- SEPs were switched to Epo-containing SEEM for another 5-day culture in normoxia (20% $O_2$). SEPs were harvested at indicated time points for qRT-PCR analysis of indicated erythroid-specific genes (n=3 per condition, unpaired t test).

Data represent mean ± SEM. n.s. $p > 0.05$, ** $p < 0.01$.
